# Inferring kinetic parameters of oscillatory gene regulation from single cell time series data

**DOI:** 10.1101/2021.05.12.443895

**Authors:** Joshua Burton, Cerys S. Manning, Magnus Rattray, Nancy Papalopulu, Jochen Kursawe

## Abstract

Gene expression dynamics, such as stochastic oscillations and aperiodic fluctuations, have been associated with cell fate changes in multiple contexts, including development and cancer. Single cell live imaging of protein expression with endogenous reporters is widely used to observe such gene expression dynamics. However, the experimental investigation of regulatory mechanisms underlying the observed dynamics is challenging, since these mechanisms include complex interactions of multiple processes, including transcription, translation, and protein degradation. Here, we present a Bayesian method to infer kinetic parameters of oscillatory gene expression regulation using an auto-negative feedback motif with delay. Specifically, we use a delay-adapted nonlinear Kalman filter within a Metropolis-adjusted Langevin algorithm to identify posterior probability distributions. Our method can be applied to time series data on gene expression from single cells and is able to infer multiple parameters simultaneously. We apply it to published data on murine neural progenitor cells and show that it outperforms alternative methods. We further analyse how parameter uncertainty depends on the duration and time resolution of an imaging experiment, to make experimental design recommendations. This work demonstrates the utility of parameter inference on time course data from single cells and enables new studies on cell fate changes and population heterogeneity.

## 1 Introduction

The identification of regulatory mechanisms that control gene expression may have important implications in biological systems. Cell state transitions are a key contributor to many processes in healthy and diseased tissue, and as such they play a major role in development, regeneration, and cancer. There is an increasing amount of literature uncovering the relationship between gene expression dynamics, i.e. dynamic changes in protein copy numbers from a single gene, and cell state transitions^[1–7]^. For example, Imayoshi et al. (2013)^[1]^ used optogenetics to show that oscillatory expression of the transcription factor ASCL1 promotes cell proliferation of mouse neural progenitor cells, whereas sustained expression promotes differentiation. Manning et al. (2019)^[2]^ linked aperiodic HES5 protein expression dynamics to murine neural progenitors, and declining oscillatory dynamics to differentiating neurons. Further evidence by Soto et. al. and Phillips et. al.^[3;8]^ demonstrates the contribution of gene expression noise to tuning oscillatory dynamics and influencing dynamically driven cell state transitions.

Experimentally, the dynamics of gene expression can be studied using a variety of approaches. Accurate measurements of protein dynamics are made through live-imaging of transcription factors in single cells, which provides real-time information on gene regulation and identifies cell-to-cell heterogeneity. This can be achieved through fluorescent fusion reporters^[9]^, where endogenously expressed proteins are attached to fluorescent reporter molecules. Fluorescence microscopy can then be used to obtain time series data that quantify protein expression levels over time (Figure 1A and B, supplementary section S.10). It may further be possible to translate the fluorescence intensity into exact protein copy numbers^[2;3]^. Fluorescent protein reporters are widely used to research the role of transcription factor dynamics in cell differentiation events, and have provided dynamic data on gene expression in various contexts, such as neural differentiation, circadian regulation, and cell cycle regulation^[1;2;10–13]^.

**Figure 1:**
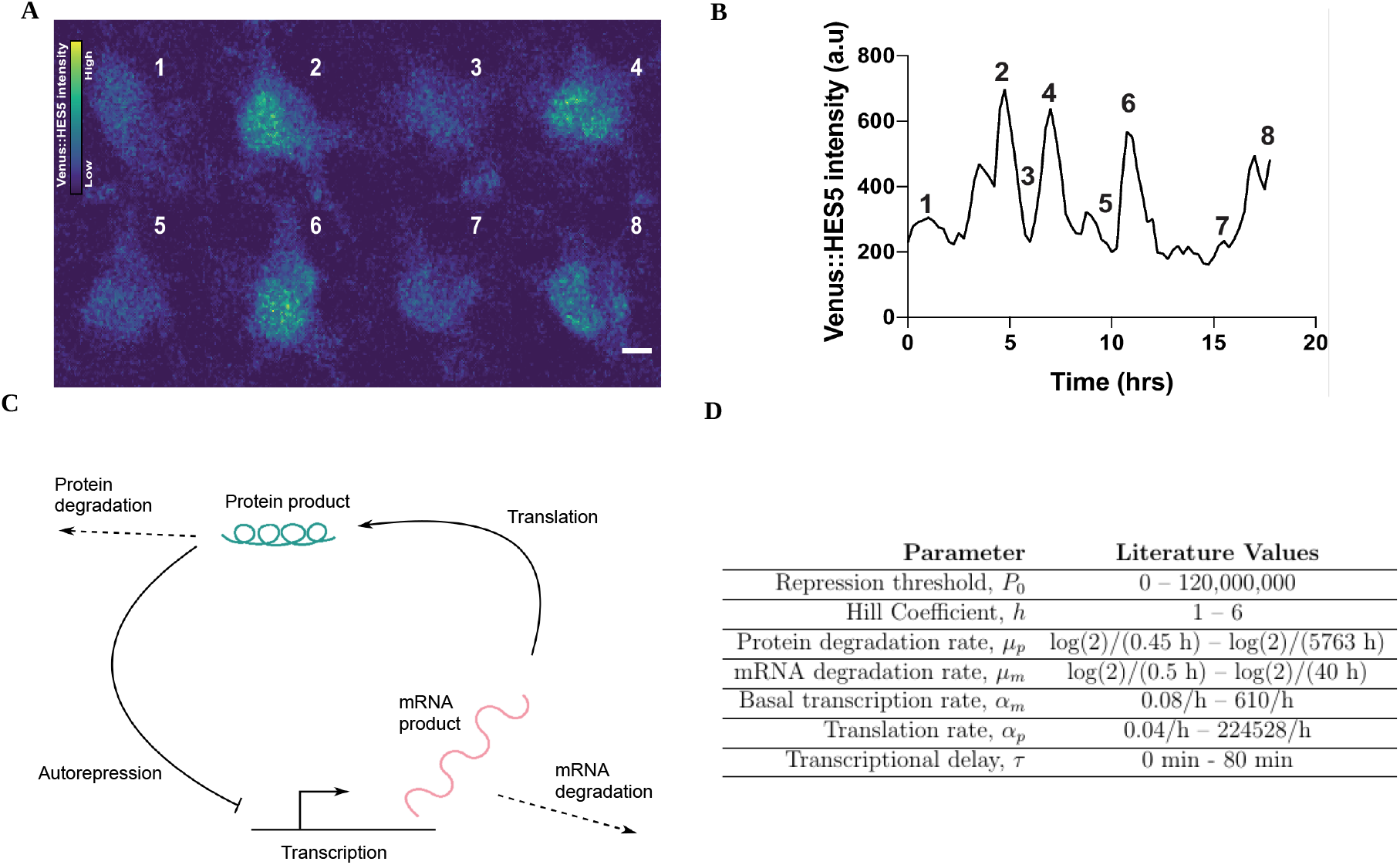
Time series data of protein expression can be modelled with an auto-negative feedback motif. **A**. Stills from a movie of a single cortical neural progenitor in vitro with Venus::HES5 knock-in reporter. Colour bar shows Venus::HES5 intensity. Stills taken at time points 1.75h (1), 4.5h (2), 6h (3), 7h (4), 9.5h (5), 10.75h, (6), 15.5h (7), 17.25h (8). Scale bar 5µm. For details on data collection see supplementary section S.10.1. **B**. Venus::HES5 intensity time series of cell in A. **C**. Graphical representation of the auto-negative feedback motif. **D**. Model parameter values taken from previously published experiments and theoretical considerations.^[2;8;19–23]^.

Mechanistically, dynamic gene expression is controlled by multiple processes, including transcriptional pulsing (transcription occurring in pulses or bursts), stochastic fluctuations (due to a limited number of molecules), gene regulatory interactions and translational control. In order to understand how these processes interact to modulate gene expression dynamics it is necessary to use mathematical models.

Within systems biology, mathematical models are often represented as a collection of gene regulatory motifs^[14;15]^. One very common motif is the delay-mediated, auto-repressive negative feedback loop (Figure 1C), which gives rise to oscillations and other dynamic patterns of gene expression that have been observed in somitogenesis, neurogenesis, and in cancer cell lines^[2;3;7;10;16]^. In this motif, a protein represses the transcription of its own gene. In combination with delays that are intrinsic to biological systems, this admits a range of dynamic behaviours, most notably oscillations at the mRNA and protein level. Regulation of gene expression through the auto-negative feedback motif contributes to cell state changes in multiple systems, including neural differentiation^[2;17;18]^.

Despite great advances in the collection of dynamic data on gene expression, and the modelling of these data, challenges remain when calibrating models to data. Even simple mathematical models, such as the auto-negative feedback motif (Figure 1C), employ multiple model parameters that correspond to biophysical quantities. For example, the auto-negative feedback motif uses rates of transcription, translation, degradation, and other parameters to predict protein and mRNA expression dynamics. Each of these parameters can take a large range of values (Figure 1D). For many application areas, parameter inference, i.e. identifying which parameters correspond to a given experimentally obtained data set, remains an open problem, since it requires the ‘inverse’ of the model, which typically cannot be computed directly. However, solving this problem bears great potential for the research of gene expression dynamics and its links to cell fate. Identifying which parameter changes correspond to observed differences in protein expression dynamics may illuminate the molecular pathways that contribute to cell fate control, and identify new sources of heterogeneity within a cell population.

The need for parameter estimation in biological systems has motivated extensive research in recent years, with a variety of approaches being developed for different types of data^[24–27]^. Techniques using Bayesian inference have emerged as a preferred approach due to their ability to quantify uncertainty in the face of noisy data, which is a common feature of biological experiments^[28]^, by representing parameters with distributions, rather than point estimates^[20;29–35]^. Placing probability distributions over our parameters, rather than treating them as point estimates, allows us not only to determine the most likely values for each of the parameters, given some data, but also to quantify our uncertainty in them.

To achieve parameter estimation with uncertainty quantification, Bayesian inference aims to identify the posterior distribution of the model under consideration, denoted *π*(***θ*** | **y**), where ***θ*** and **y** are the model parameters and observed data respectively. The posterior distribution describes the probability of the model parameters given observed data, and can be calculated using Bayes’ rule

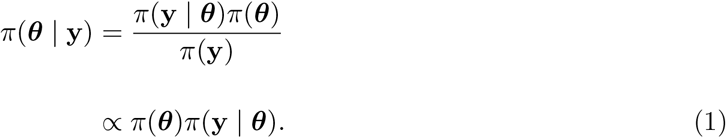

Here, *π*(**y** | ***θ***) is referred to as the likelihood, and is a measure of the fit of a statistical model to the observed data, given specific values of the model parameters. The prior probability, *π*(***θ***), is a distribution which outlines one’s beliefs in the parameters ***θ*** before any new data is taken into account. These prior distributions can be informed using published data (Figure 1D), as well as physical constraints (e.g. rate constants must be positive). To visualise the posterior distribution and use it in further analysis it is common to work with computationally generated samples from this distribution. Posterior probabilities may be difficult to compute directly, hindering the efficient generation of these samples^[36;37]^.

Specifically, it may not be possible to calculate posterior probabilities if the likelihood of the model is not available. In these cases, Approximate Bayesian Computation (ABC) can be used. However, ABC reduces the data to a small number of summary statistics, which inevitably decreases the accuracy of inference^[38]^. If an expression for the likelihood is available and can be calculated at given parameter points, the calculation of the marginal likelihood *π*(**y**) often poses a further challenge in Bayesian inference, since it may require the numerical integration of the likelihood and prior probability. To overcome this challenge, sampling from the exact posterior distribution can be achieved using Markov chain Monte Carlo (MCMC) techniques, such as the Metropolis-Hastings random walk^[39]^.

MCMC methods can produce samples from the posterior distribution *π*(***θ*** | **y**) even if the integration factor *π*(**y**) is unknown. In many scenarios, the reconstruction of a posterior distribution using MCMC sampling can be slow, in particular if the parameter space is high-dimensional, if the calculations of the likelihood are computationally expensive, or if parameters are highly correlated within the posterior distribution^[40]^. In these scenarios, more efficient Hamiltonian Monte Carlo (HMC) or Metropolis-adjusted Langevin algorithm (MALA) methods are preferable^[41–43]^. HMC and MALA algorithms additionally require the gradient of the posterior probability with respect to the model parameters and can result in orders-of-magnitude faster convergence of the sampled distribution to the posterior distribution, especially for high-dimensional distributions or when parameter correlations are present^[42;44]^.

For time series data specifically, a common approach to calculating the likelihood is the Kalman filter. The Kalman filter is an algorithm which calculates the likelihood of the data at each time point, given a mathematical model of stochastic dynamics, and an observation noise model. It can generally be applied to Markov processes, where dynamic changes over time only depend on the current state of the system, and not past states. The Kalman filter is a powerful method to calculate posterior probabilities if delays are not present in the model^[45]^, and can be extended to estimate the gradient of the likelihood function, making gradient-based sampling of the posterior distribution possible^[46]^.

A number of recent methods focus specifically on time series of gene expression^[2;26;47–51]^. For the study of oscillatory gene expression, a wide array of studies discuss time series data of protein concentrations, such as in Figure 1A,B, as well as the description of these data through the auto-negative feedback motif (Figure 1C). Despite this, a reliable Bayesian inference method for this popular combination of data and model is still missing. Since the model includes delays, the widely-used Kalman filter approaches are not applicable. Recently, Calderazzo et al. (2018) have addressed this problem by identifying a method to introduce delays into the Kalman filter^[52]^, indicating that accurate Bayesian inference for the auto-negative feedback motif on time series data of gene expression may be possible. However, this approach lacks the ability to calculate gradients of the posterior probability distribution, thus preventing the use of efficient gradient-based sampling methods. Furthermore, while Calderazzo et al. (2018) applied their method to a motif containing negative feedback, this method has not yet been applied to the widely used motif in Figure 1C, which includes mRNA in addition to protein.

Here, we present a Bayesian inference pipeline that can be used as a non-invasive method to measure kinetic parameters of gene expression emerging from the auto-negative feedback motif using protein expression time course data. We extend the Kalman filtering method presented by Calderazzo et al. (2018) by introducing a recursive implementation to calculate the gradient of the likelihood. This enables us to embed the non-linear delay-adapted Kalman filter into a state-of-the-art MALA sampling algorithm. This extension enhances the robustness of the inference, making it more suitable for use in typical experimental settings.

Our method is able to capture multiple kinetic parameters of gene expression simultaneously using time course data from single cells, and outperforms previous approaches. We demonstrate the accuracy of our method on *in silico* data, provide an example on how the method can be applied to experimental data, and show how the method can be used to obtain experimental design recommendations. This work is paving the way for the use of Bayesian inference methods for the investigation of gene expression dynamics and their links to cell fate.

## 2 Methods

In this section we give an overview of the key components of our method. First, we introduce the mathematical model for the auto-negative feedback motif. Then, we discuss how we use a delay-adapted non-linear Kalman filter to approximate the likelihood function. Lastly, we provide details on data processing. Descriptions of our method that require longer derivations, as well as further details on data collection, are provided in the Supplementary Information. This includes our implementation of two MCMC methods, Metropolis-Hastings random walk (MH) and MALA, as well as our proposed algorithm to compute the gradient of the likelihood function, which is a major technical advancement in this paper. The availability of this gradient enables the use of a wider range of MCMC samplers, such as MALA, which we use throughout the paper.

### 2.1 The negative feedback chemical reaction network

Here we consider a widely used model of gene expression, that incorporates knowledge of the auto-repressive negative feedback loop (Figure 1C). Our model describes both protein and mRNA expression dynamics over time at the level of a single cell, accounting for transcription and translation, as well as degradation. We include a delay in the model, representing the time taken from the initiation of transcription until the production of a transcript and its removal from the nucleus. We further account for the effect of transcriptional auto-repression, where a high abundance of the target protein inhibits transcription of the mRNA^[22;23;53]^.

Let *p*(*t*) and *m*(*t*) define the number of protein and mRNA molecules, respectively, at time *t* for a gene of interest. Gene expression is often subject to stochastic effects due to finite molecule numbers. To reflect this we model the system with delayed Chemical Langevin Equations^[54–56]^,

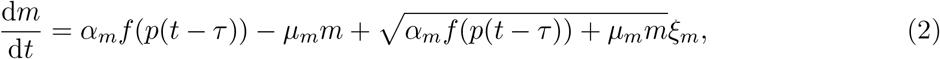

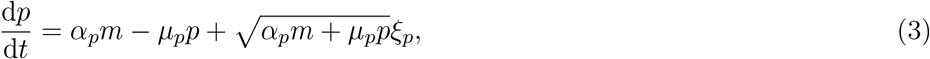

where *ξ*_*m*_, *ξ*_*p*_ denote Gaussian white noise, i.e.

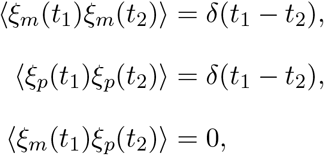

where *δ*(·) is the Dirac-delta function.

The parameters *µ*_*m*_, *µ*_*p*_, *α*_*m*_, and *α*_*p*_ describe the rate of mRNA degradation, protein degradation, basal transcription rate in the absence of protein, and translation rate respectively. The transcriptional delay is given by *τ*, and auto-repression is taken into account via the use of a Hill function

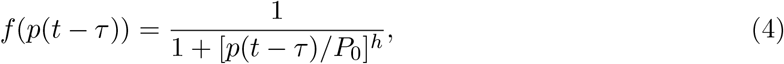

reducing the rate of transcription for increasing amounts of protein *p* at time *t* − *τ*^[57]^. Here, *τ*, the time delay, is the duration of the transcription process. The Hill function (eq. (4)) is close to one when the protein at time *t* − *τ* is much less than the repression threshold *P*_0_ and close to zero when the the protein at time *t* − *τ* is much more than the repression threshold. The steepness of the transition from one to zero can be regulated by the Hill coefficient *h*. The Hill coefficient reflects the extent of cooperativity between ligand binding sites for the gene of interest^[58]^.

From equations (2) and (3) we can see that the instantaneous rate of transcription is determined by *α*_*m*_*f* (*p*(*t* − *τ*)). This allows us to define an approximation for the average rate of transcription as

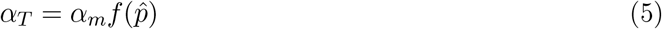

Where 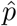 is the average expression of protein. This average expression of protein may be obtained from simulated or experimental data.

We simulate the stochastic differential equations (2) and (3) using the Euler-Maruyama method with a time step Δ*t* = 1 min, which is chosen sufficiently small to ensure numerical accuracy of the scheme.

A deterministic version of the model in equations (2) and (3) was first developed by Monk (2003) in order to describe gene expression oscillations of Hes1, p53 and NF-*κ*B, and various versions of the model have since been widely studied^[2;8;21–23;55;56]^. In particular, when molecular copy numbers of mRNA and protein are low, we expect the rate processes of transcription, translation, and degradation to stochastically vary with time. This effect is accounted for by the noise terms in the Chemical Langevin Equations (2) and (3)^[54–56]^.

### 2.2 The likelihood function can be evaluated through Kalman filtering

The Kalman filter is an algorithm which calculates the likelihood function for linear stochastic differential equations describing time-series data^[59]^. The Kalman filter evaluates the likelihood of each time-point recording consecutively. The full likelihood is then the product of these individual likelihoods, exploiting the Markov property of the underlying stochastic process. The Kalman filter can be extended to non-linear dynamical systems by using piecewise-linear Gaussian approximations^[60]^.

Here, we implement a Kalman filter, extended to account for non-linearity and delay, in order to evaluate the likelihood that our observed data results from the model in eqs. (2) and (3) at a given parameter combination. This likelihood can then be used to infer model parameters for a given experimentally observed time series recording of gene expression. The resulting posterior distribution may then represent testable predictions on the biophysical characteristics of the gene of interest, such as transcription, translation, and degradation.

Our Kalman filter implementation uses a finer discretisation on the time axis than that given by the observation interval. Specifically, we introduce *z* hidden states between consecutive observations. Introducing such hidden states is common when applying Kalman filters to non-linear stochastic differential equations. It increases the accuracy of a piece-wise linear Gaussian approximation. In the following, the time variable *t* will assume integer values numbering all discretisation time points, i.e. *t* = 0, 1, …, *nz*, where *n* is the total number of observations.

It is possible to show that the likelihood of a set of observations given specific model parameters can be expressed as^[52]^

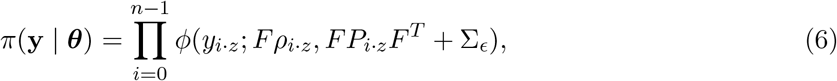

where the subscript *i* · *z* denotes multiplication of *i* and *z* and

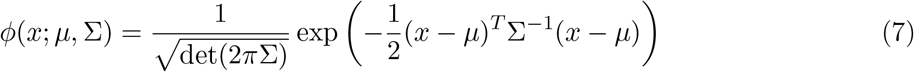

is the multivariate Normal distribution. The true, unobserved state of the system at time *t* is given by ***X***(*t*) = *x*_*t*_ = [*m*(*t*), *p*(*t*)]^T^, and the relationship between *x*_*t*_ and the observed data *y*_*t*_ is given by *y*_*t*_ = *Fx*_*t*_ +*∈*_*t*_, where *∈*_*t*_ ∼ 𝒩 (0, Σ_*ϵ*_) and *F* is a 1 × 2 matrix. Thus, *F* and represent our measurement model. Throughout, we use *F* = [0, 1], since we aim to apply our method to data on protein expression dynamics, where measurements of mRNA levels are not available. The value Σ is called the measurement variance, and describes the observation noise introduced through the experimental measurement process. The variables *ρ* and *P* represent the *state space mean* and *state space variance* respectively. We define *y*_0:*t*_ = [*y*_0_, *y*_*z*_, *y*_2*z*_, …, *y*_*t*_]^*T*^, and write *ρ*_*t*_ = E[**X**(*t*) | *y*_0:*t−*1_] and *P*_*t*_ = Cov(**X**(*t*), **X**(*t*) | *y*_0:*t−*1_).

The Kalman filter calculates *ρ*_*t*_, and *P*_*t*_ in eq. (6) using an iterative process with two main steps. At iteration *k*, the first *k* observations have been used to infer a probability distribution over the true state of the system **X**(*t*) for all discretisation time points up to *t* = *kz*. This probability distribution is characterised by it’s mean 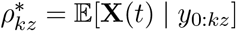 and covariance 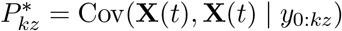.

In the Kalman filter *prediction step* we then use the model to calculate the predicted probability distribution for protein and mRNA copy numbers at the next observation time point, **X**((*k* + 1)*z*). We use this prediction to evaluate the likelihood of the observed data at the *k* + 1 observation time point. Before the prediction for the next observation is made, the Kalman filter *update step* is applied, in which the probability distribution of the state space up to observation *k* + 1 is updated to take the measurement at *t* = (*k* + 1)*z* into account.

For our update step we derive an expression for the mean and variance of the state space distribution *π*(*x*_*t−τ*:*t*_ | *y*_0:*t*_), denoted 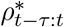 and 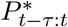 respectively. That is, the likelihood of our state space estimates from the past time *t* − *τ* to the current time, *t*, given all of our current observations. This is necessary in order to accurately predict the state space distribution at the next observation time point, *π*(*x*_*t*+Δ*t*_ | *y*_0:*t*_), as past states can affect future states due to the presence of delays. We provide detailed derivations of our Kalman filter prediction and update steps in the Supplementary Information Section S.1.

### 2.3 Implementation of MCMC sampling algorithms

The aim of our inference algorithm is to generate independent samples from the posterior distribution, *π*(***θ*** | **y**). In this paper, we compare results from two different sampling methods, MH and MALA. The MH algorithm and MALA are two of the most widely used MCMC methods for drawing random samples from a probability distribution. For completeness, we provide their algorithms in the Supplementary Information Sections S.2 and S.3.

Drawing proposals using MALA requires the calculation of the gradient of the log-posterior *U* (***θ***), which we outline in Section S.4. This is achieved by iteratively computing the derivatives of state space mean, *ρ*_*t*_, and state space variance, *P*_*t*_, with respect to each parameter, as detailed in Section S.5.

### 2.4 Trends in the data are identified by Gaussian processes

Before applying our inference method we detrend protein expression time series using Gaussian process regression, in order to identify and exclude data that show significant long-term trends^[61;62]^ (see Section 3.3 for further motivation). Specifically, we make use of a *scaled* squared exponential Gaussian process combined with white noise, whose kernel is given by

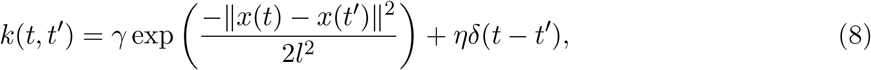

where *x*(*t*) − *x*(*t*′) is the Euclidean distance between *x*(*t*) and *x*(*t*′), *l* is the lengthscale, and *γ, η* ∈ (0, ∞). In the Gaussian process regression the hyperparameters *γ, l*, and *η* are found using constrained optimisation.

The initial value of the lengthscale is 1000 minutes, and is bounded uniformly in the range (1000 min, 2000 min). The lower bound of this range, 1000 minutes, was chosen to ensure that detrending does not perturb ultradian dynamics in the data. The upper bound, 2000 minutes, was chosen sufficiently large to ensure that detrending is not affected by it. The initial value of the parameter *γ* is the variance of the data, 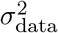, and is restricted by a uniform prior to 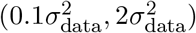. The parameter *η* has initial value 100, and is restricted by a uniform prior to 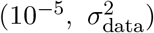. Here, *x*(*t*) and *x*(*t*′) represent our protein expression time course data at time *t* and *t*′ respectively. We identified data without a significant long term trend manually by visual inspection (see Section 3.3, Figure 4) and removed any residual trend before applying our inference method.

## 3 Results

Single cells in a seemingly homogeneous population can change cell fate based on gene expression dynamics. The control of gene expression dynamics can be understood with the help of mathematical models, and by fitting these models to experimentally measured data. Here, we analyse our new method for parameter inference on single-cell time series data of gene expression using the widely used auto-negative feedback motif. We first validate our method by showing the performance of our algorithm on *in silico* data sets. We then demonstrate the utility of our method by applying it to experimentally measured data and, finally, use our method to analyse how parameter uncertainty may depend on properties of the data, as well as the experimental design.

### 3.1 Sampled posterior distributions agree with analytical derivations for one-dimensional parameter inference

We first test our inference method on *in silico* data from the forward model of the auto-negative feedback motif (Figure 1C). This is done using Chemical Langevin Equations, as detailed in Section 2.1. Specifically, we emulate an in silico imaging experiment by selecting simulated data in sparse intervals of Δ*t*_obs_ mins and mimic measurement noise by adding random perturbations to each observation time point (Figure 2A). These perturbations are drawn from a Gaussian distribution with variance Σ_*ϵ*_. Testing the method on *in silico* data first is beneficial, since ground truth parameter values are known *a priori* for the generated *in silico* data sets, and can be compared to the obtained posterior distributions.

**Figure 2:**
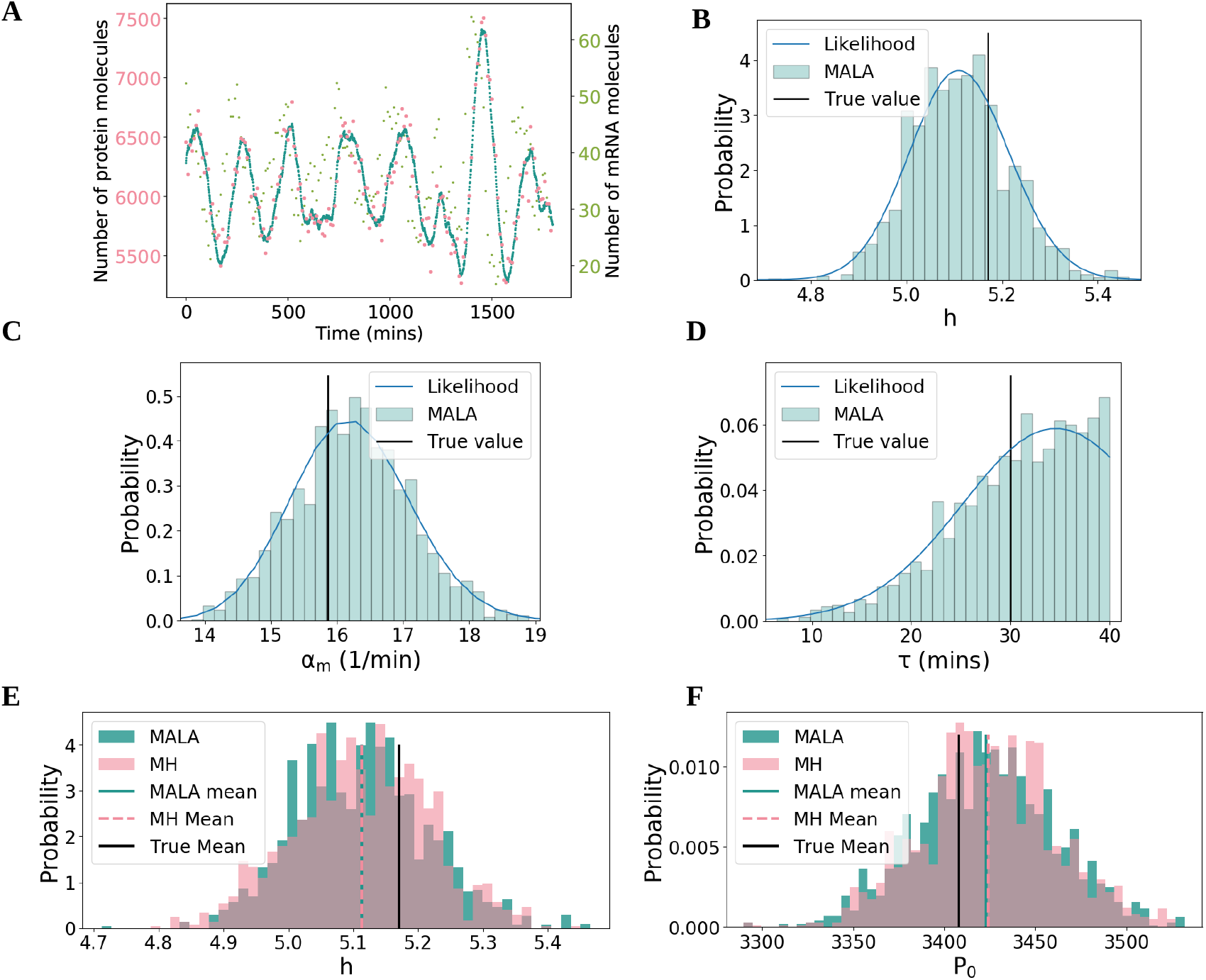
Our algorithm accurately samples posterior distributions. **A**. Simulated experimental data. Protein copy numbers are simulated using the Chemical Langevin Equation (see Section 2, blue dots). Experimental observations are emulated every five minutes by adding Gaussian noise to the protein copy number (pink). The parameter values used were *P*_0_ = 3407.99, *h* = 5.17, *µ*_*m*_ = log(2)*/*30, *µ*_*p*_ = log(2)*/*90, *α*_*m*_ = 15.86, *α*_*p*_ = 1.27, *τ* = 30, Σ = 10000, and the simulated mRNA copy numbers are also included (green dots). **B, C, D**. Posterior distributions for one-dimensional inference. For individual model parameters, posterior distributions were inferred while keeping all other parameters fixed, respectively. Shown above are the inferred marginal posteriors for the Hill coefficient (B), transcription rate (C), and transcriptional delay (D) respectively as histograms, using MALA as the underlying sampling algorithm (see Section S.2) for 2500 iterations. The blue lines are the exact likelihood calculations. The sampled and exact distributions coincide. **E, F**. Histograms for both MALA and MH on the 1-dimensional problem for the Hill coefficient (E) and repression threshold (F).

We start by applying our inference method to simple test cases, where the true values of all but one parameter are known, and only the remaining, unknown, parameter value is inferred (Figure 2). This allows us to compare our sampled posterior distributions to the exact likelihood, which can be calculated in these one-dimensional examples using equation eq. (6). If our inference method is accurate, the sampled posterior distribution should closely match the exact likelihood if the Markov chain has converged (see Section S.7). We find that this is indeed the case for example *in silico* data sets (Hill coefficient, transcription rate and transcriptional delay in Figure 2B–D, repression threshold and translation rate in Figure S1). Additionally, ground truth parameter values lie well within the support of the posterior distribution (Figure 2B–D, supplementary Figure S1, vertical black lines).

Our proposed inference method uses the MALA sampler, which relies on calculating likelihood gradients (see Section S.4). The comparison with exact calculations in Figure 2B–D and supplementary Figure S1 validates our implementation of MALA, and the associated computations of the likelihood gradient. In order to further test our implementation of MALA, and the associated computations of the likelihood gradients, we compare our results to posterior distributions sampled using the MH algorithm, which does not require gradient calculations. Despite an expected slower convergence of the MH algorithm, this comparison is feasible for one-dimensional posterior distributions, which typically can be well approximated with a few thousand samples. The sampled means have a relative difference below 0.03%, and the standard deviations fall within 4% of each other (Table 1, supplementary Table S3). This comparison reveals that posterior distributions from both samplers agree well with each other (Figure 2E–F, Figure S2), and further validates the implementation of the individual likelihood gradients.

**Table 1:** The true values for the parameters which were used to generate the data in Figure 3A, alongside the means, ***µ***, and standard deviations, ***σ*** of the corresponding one-dimensional posterior distributions, from both the MALA and MH algorithms (Figure 2E,F).

| Parameter | True Value | $\mu$ (MALA) | $\mu$ (MH) | $\sigma$ (MALA) | $\sigma$ (MH) |
| --- | --- | --- | --- | --- | --- |
| Repression threshold, $P_0$ | 3408 | 3422 | 3423 | 37.51 | 36.65 |
| Hill Coefficient, $h$ | 5.17 | 5.113 | 5.112 | 0.100 | 0.104 |

### 3.2 Our method allows for simultaneous inference of multiple model parameters

Having validated the method on one-dimensional posterior distributions, we further test the performance of the method by simultaneously inferring multiple model parameters from a single *in silico* data set and comparing the resulting posterior distribution to the ground truth parameter combination (Figure 3A,B). Since we cannot measure convergence of the sampled posterior through comparison to the true posterior distribution in the multi-dimensional case, we rely on typical MCMC convergence diagnostics (Section S.7).

**Figure 3:**
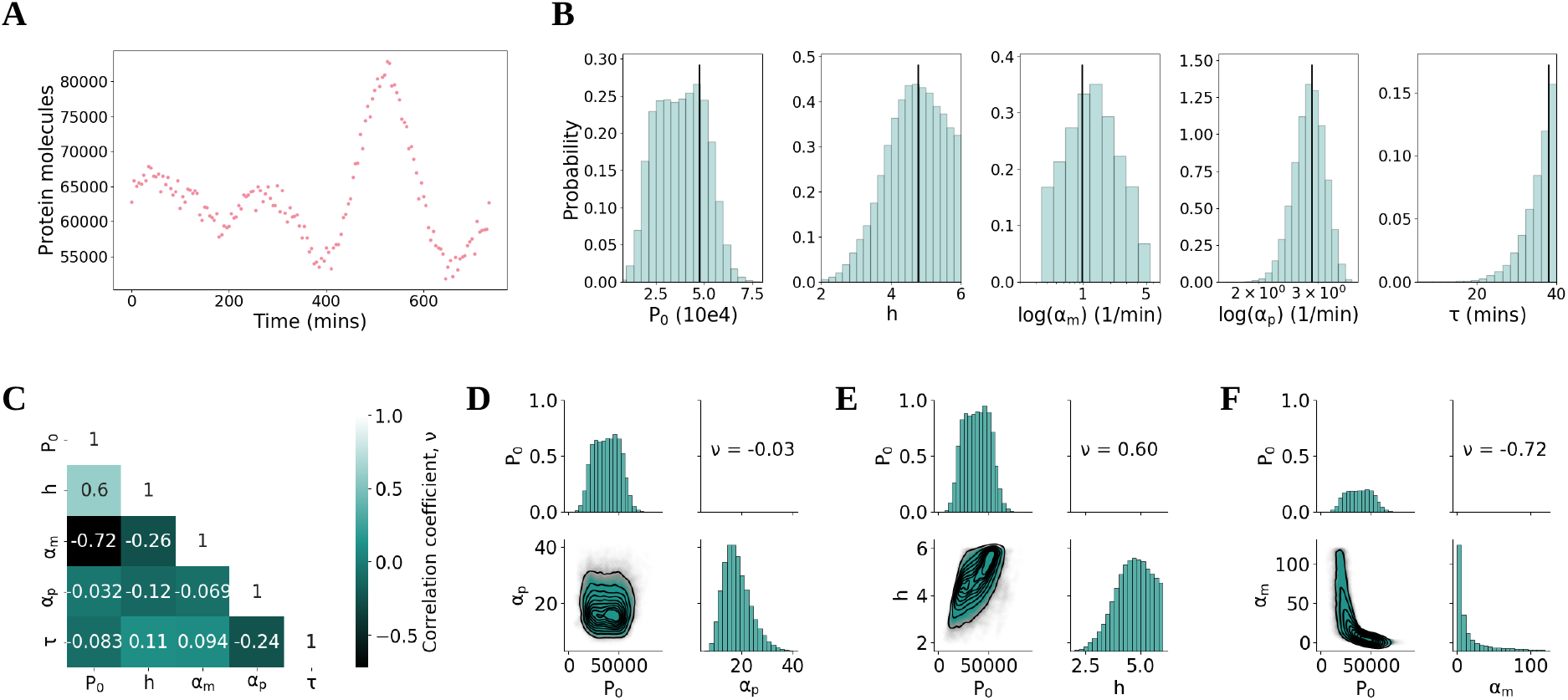
MCMC sampling enables simultaneous inference of multiple parameters. **A**. An *in silico* data set was generated using parameter values *P*_0_ = 47515, *h* = 4.77, *µ*_*m*_ = log(2)*/*30, *µ*_*p*_ = log(2)*/*90, *α*_*m*_ = 2.65, *α*_*p*_ = 17.61, *τ* = 38.0, Σ_*ϵ*_ = 10^6^. **B**. Our method is applied to the data set in (A) to sample the joint posterior distribution over five parameters. The marginal posteriors for each parameter are shown. All marginal posterior means are within half a standard deviation of the true value. **C**. A (symmetric) correlation matrix which shows the correlation coefficient, *ν*, between samples for each pair of parameters. Diagonal entries are, by definition, perfectly correlated (*ν* = 1), and off diagonal entries take values in the range [−1, 1]. **D,E,F**. Joint posterior distributions showing the relationship between different pairs of parameters. D shows the repression threshold, *P*_0_ is not correlated with the protein translation rate, *α*_*p*_. E shows a correlation of *ν* = 0.6 between the repression threshold and the Hill coefficient, *h*. In other words, for this data set, a higher Hill coefficient indicates a higher repression threshold (and vice-versa). Finally, F shows the strong negative correlation between *P*_0_ and *α*_*m*_.

We choose a data set that shares characteristics with typically collected time course data from single cells. Specifically, our *in silico* data set is of similar length and observation intervals as previously analysed by Manning et al. (2019). In this paper, the degradation rates of protein and mRNA have been measured, so we assume these measurements as known values, leaving five unknown parameter values to infer. The prior distributions were uniform, defined by the range of values given in Table S1, and log-uniform for *α*_*m*_ and *α*_*p*_ (see Section S.6 for details).

We find that the marginal posterior means, i.e. values of largest probability, all lie within maximally half a standard deviation of the ground truth values (Table 2). This indicates that a high degree of accuracy in the inference can be achieved with the amount of data typically gathered from a single cell.

**Table 2:** The true values for the parameters which were used to generate the data in Figure 2A, alongside the means, ***µ***, and standard deviations, ***σ***, using MALA.

| Parameter | True Value | Mode | SD |
| --- | --- | --- | --- |
| Repression threshold, $P_0$ | 47515 | 44915 | 12434 |
| Hill Coefficient, $h$ | 4.77 | 4.41 | 0.80 |
| Log basal transcription rate, $\log(\alpha_m)$ | 0.975 | 1.45 | 1.24 |
| Log translation rate, $\log(\alpha_p)$ | 2.869 | 2.808 | 0.288 |
| Transcriptional delay, $\tau$ | 38.0 | 39.87 | 4.39 |

Simultaneous inference of multiple parameters further allows for the investigation of pairwise parameter correlations, using correlation coefficient *ν* (Figure 3C). Pairwise correlations provide crucial information on how posterior distributions can be constrained further. Specifically, the strong correlation between the repression threshold, *P*_0_, and the logarithm of the basal transcription rate, log(*α*_*m*_) (Figure 3E), highlights that the data in Figure 3A is consistent with either high repression thresholds and low transcription, or vice versa. Such strong pairwise correlations (Figure 3E,F) imply that gaining new information on one of the two parameters would constrain the other. This is not the case when parameters are uncorrelated, such as the transcriptional delay and the translation rate (Figure 3D), and experimentally measured values on either of these parameters would not inform the other.

### 3.3 Parameter inference on single cell data outperforms previous approaches and may reveal underlying mechanisms for population heterogeneity

Next, we seek to evaluate the performance and utility of our method by applying it to experimentally measured data. Specifically, we investigate data on gene expression oscillations in mouse spinal cord neural progenitor cells^[2]^ (see supplementary section S.10.2), and compare our method to results on parameter inference from ABC (Figure 4A). In this previous approach, inference was performed using population-level summary statistics of the collected time-course data. This resulted in posterior distributions with high parameter uncertainty. Specifically, the marginal posterior distributions for the Hill coefficient and the transcriptional delay were close to uniform, illustrating that the provided summary statistics did not contain sufficient information to constrain the uniform prior distribution. The remaining parameters had distinct modes. Nonetheless, parameter uncertainty was high since the spread of the posterior distribution was comparable to that of the prior^[2]^. Importantly, this previous approach did not allow for comparison of posterior distributions between single cells. When applying our method to time series data from fluorescence microscopy experiments, it is necessary to address that our model of the auto-negative feedback motif cannot describe longterm trends in data. Specifically, the model of the auto-negative feedback loop considered here is designed to describe ultradian oscillations that typically have periods shorter than 10 hours^[21;22;55]^, and cannot describe variations in protein numbers on longer timescales, such as one would expect from a slow up- or down-regulation of the gene in the tissue. Hence, we only apply our algorithm to protein expression time series that we expect to be accurately modelled by eqs. (2) and (3) by excluding data that show significant long-term trends. In order to identify such time series, we first remove trends from the time series that vary on lengthscales longer than 10 hours by using Gaussian process regression (see Section 2.4). Then, we manually identify all time series for which the detrended and raw time series visually agree (Figure 4B) and select these for inference.

**Figure 4:**
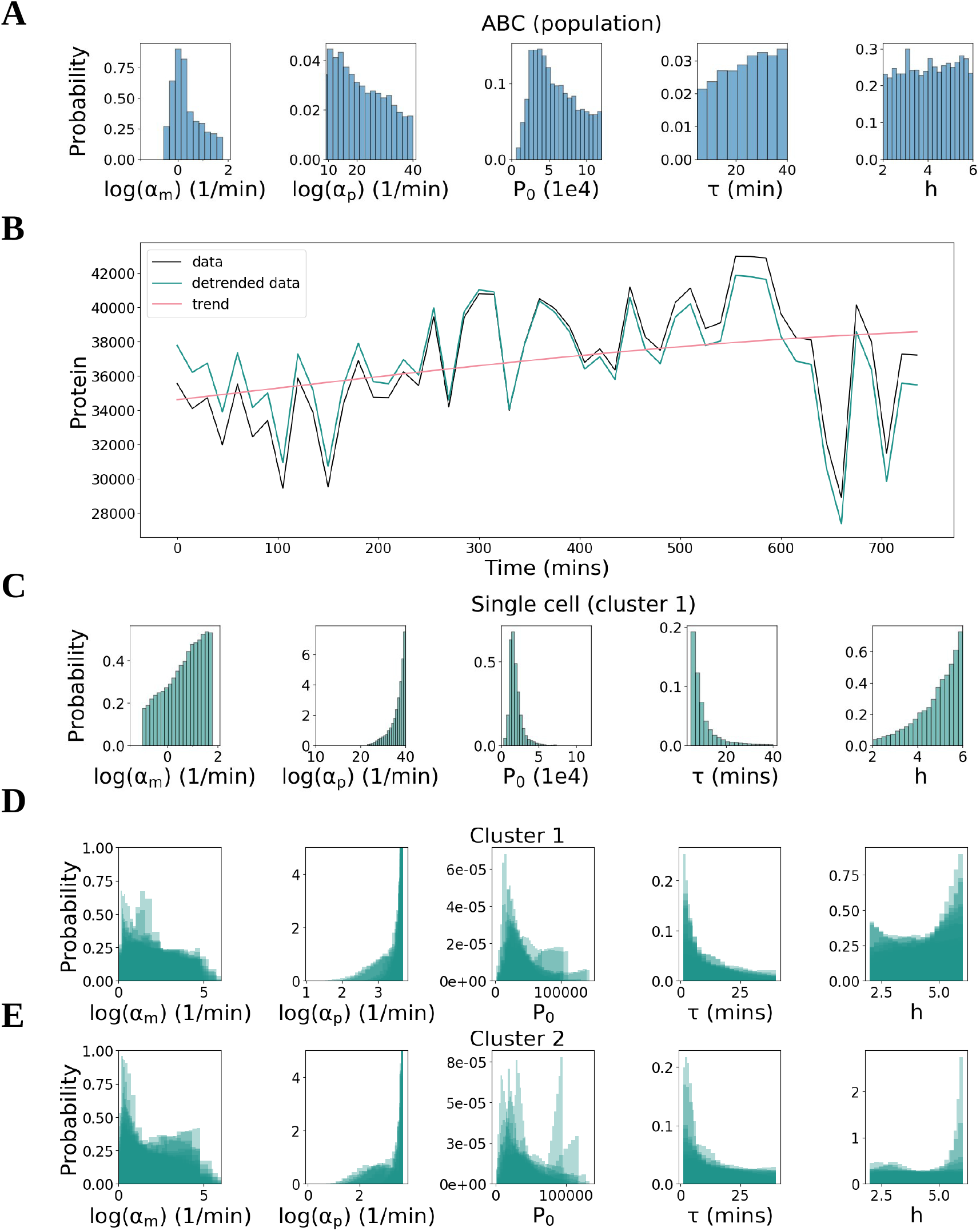
Parameter inference on single cell data outperforms previous approaches. **A**. Marginal posterior distributions for each parameter are obtained from the ABC algorithm using summary statistics in place of likelihood evaluations. **B**. Detrending of *in vivo* single cell protein expression data (black line). The sampling interval is 15 minutes, and the length of the time course is 12 hours (720 mins). The mean is subtracted from the data and a squared exponential Gaussian process determines the long term trend (pink line). This trend is removed from the data and the mean is added back in (green line). **C**. Marginal posteriors from MALA for the detrended single cell data shown in panel B. **D,E**. Marginal posterior distributions for every cell in cluster 1 and 2 respectively.

In order to identify a suitable value for the measurement variance Σϵ we rely on previous estimates^[2]^. Manning et al. (2019) decomposed the measured time series into two contributions, one from a time-varying signal with finite auto-correlation time, and one from a time-varying signal for which consecutive observations are uncorrelated^[2]^. This second contribution follows an identical distribution as the measurement error in our model, and was estimated to contribute 10% of the total variance across all detrended time series. Hence, we set

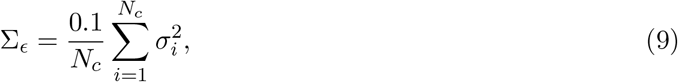

where *N*_*c*_ is the number of considered traces, and 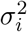 is the variance for the *i*th detrended data set.

We find that our method can identify more accurate posterior distributions than the previous ABC-based approach by using single cell time series of gene expression only (Figure 4C vs. Figure 4A.). For the single-cell gene expression time course in Figure 4B, we find that there is still comparatively high uncertainty on the basal transcription rate (*α*_*m*_ in Figure 4C), as the support of the marginal posterior distribution reflects that of the uniform prior distribution. However, for all other model parameters that are inferred from this time course, the marginal posterior distributions are narrower than the prior, and than previously identified marginal posterior distributions from ABC (Figure 4C).

Having investigated marginal posterior distributions from a single cell we proceed to analyse to what extent these posterior distributions can vary across multiple cells in the population. Among the experimental data considered here, hierarchical clustering has previously identified two sub-populations (denoted as clusters 1 and 2) which have distinct gene expression dynamics and which also do not have strong long-term trends^[2]^, such as down-regulation of gene expression. For clusters 1 and 2 there are 19 and 22 cells respectively which we have selected for negligible trends (see Section 2.4).

We find that the posterior distributions inferred from multiple cells share features that are conserved across all cells and both populations (Figure 4D,E). Specifically, the marginal posterior distributions of the translation rate *α*_*p*_ are all larger than exp(2)*/*min, and biased to larger values. Similarly, the marginal posterior distributions for the delay *τ* cover the entire range of considered values, and are biased towards smaller values, with most likely values below 10 minutes. These observations appear to hold true for both clusters considered here, and they highlight that parameter inferences obtained from our method are biologically reproducible, which is a necessary feature to enable the use of the method in practical applications.

In contrast, for the basal transcription rates *α*_*m*_ and the Hill coefficient *h*, marginal posterior distributions vary between individual cells, suggesting that there is an underlying heterogeneity of these parameters across the cell population. However, the remaining parameter uncertainty is too high to reliably identify differences between cells and clusters of cells, raising the question of how imaging protocols may need to be changed in order to achieve lower uncertainty on typical parameter estimates.

### 3.4 Longer time course data improves accuracy of inference more effectively than more frequent sampling

Typically, longer imaging time series can only be collected at the cost of a lower imaging frequency. When designing experiments, it may be desirable to choose an imaging protocol that optimises the parameter inference towards high accuracy and low uncertainty. However, parameter uncertainty may not only be influenced by the imaging protocol, but also by the bifurcation structure of the underlying dynamical system^[63]^. Hence, we next analyse how posterior distributions depend on the frequency of sampling, on the length of the imaging interval, and on the position in parameter space. To evaluate the performance of our inference, we investigate the uncertainty using *relative uncertainty*, RU_***θ***_ (Supplementary S.7, equation (S35)), which quantifies the spread of the posterior distribution. We use this metric to quantify the performance of our inference method on a number of synthetic data sets with different lengths and sampling frequencies, and for different locations in parameter space.

We choose two locations in parameter space that correspond to two different values of oscillation coherence, thus producing qualitatively different expression dynamics (Figure 5A, Table S2). The oscillation coherence is a measure for the quality of observed oscillations (supplementary Section S.8). Choosing parameter combinations with different coherence thus ensures that these correspond to different positions within the bifurcation structure of the auto-negative feedback loop^[55;64;65]^.

**Figure 5:**
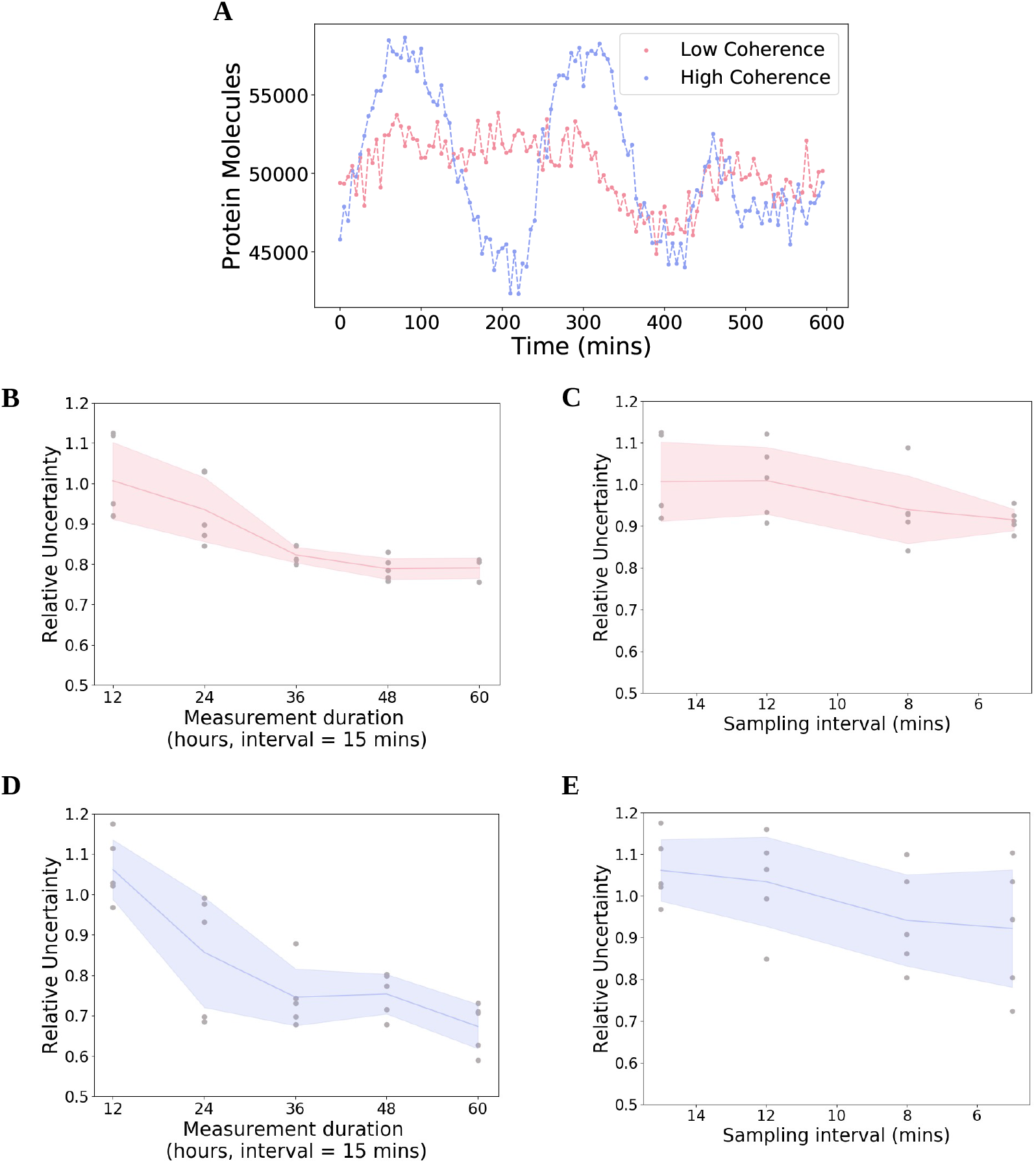
Increasing the length of time course data improves inference more than increased sampling frequency. **A**. Two examples of *in silico* protein observations, one which has low coherence (pink) and another with high coherence (purple). Exact parameter combinations can be found in Table S2. **B**. RU_***θ***_ for low coherence data sets sampled with different lengths, from 12 hours to 60 hours. As length increases from 12 hours to 36 hours, both the mean and standard deviation decrease. **C**. RU_***θ***_ for low coherence data sets sampled with different frequencies. As frequency increases from 15 minutes to 5 minutes, the mean decreases. **D**. Same as B with the high coherence data sets. **E**. Same as C with the high coherence data sets.

We first analyse to what extent collecting data for a longer sampling duration may reduce parameter uncertainty (Figure 5B,D). We find that a longer sampling duration can strongly decrease parameter uncertainty. Doubling the length of the time series reduces the uncertainty by 19% on average for the high coherence parameter combination, and 7.1% on average for the low coherence parameter combination. A tripling of the available data leads to reductions in uncertainty by 29.8% and 18.3% and for high and low coherence respectively.

In contrast, an increase in sampling frequency leads to a smaller decrease in parameter uncertainty on average (Figure 5C,E). Specifically, doubling the amount of data only leads to a decrease by 11.3% in the case of the high coherence parameter combination, and 6.7% in the case of low coherence. A tripling of the available data lead to reductions in uncertainty of 13.2% and 9.1% for low and high coherence respectively.

We find that analogous conclusions hold true if inference accuracy is analysed (ME_***θ***_, supplementary Section S.7, equation (S36)), instead of uncertainty (Figure S3). Accuracy increases with longer sampling durations and shorter imaging intervals, and longer sampling durations have a stronger effect than shorter imaging intervals.

### 3.5 Additional measurements of mRNA copy numbers improve estimates of the average transcription rate

In the previous section, we have analysed the impact of changes in the imaging protocol on parameter uncertainty overall. Alternatively, it may be desirable to identify interventions that reduce uncertainty for particular parameters of interest. For example, an important quantity of interest may be the average rate of transcription of the investigated gene, introduced as *α*_*T*_ in equation (5). In many *in silico* examples of our parameter inference, this average rate of transcription *α*_*T*_ is poorly inferred, with the mean of the posterior distribution being up to 5 times larger than the ground truth value. This is for example the case in Figure 6A. In this and other examples, the ground truth value lies outside the 85% highest density interval (HDI) of the posterior distribution (Figure 6A–C). Intuitively, one may assume that estimates for the rate of transcription are improved if measurements of mRNA copy numbers, in addition to protein expression dynamics, are considered in the inference.

**Figure 6:**
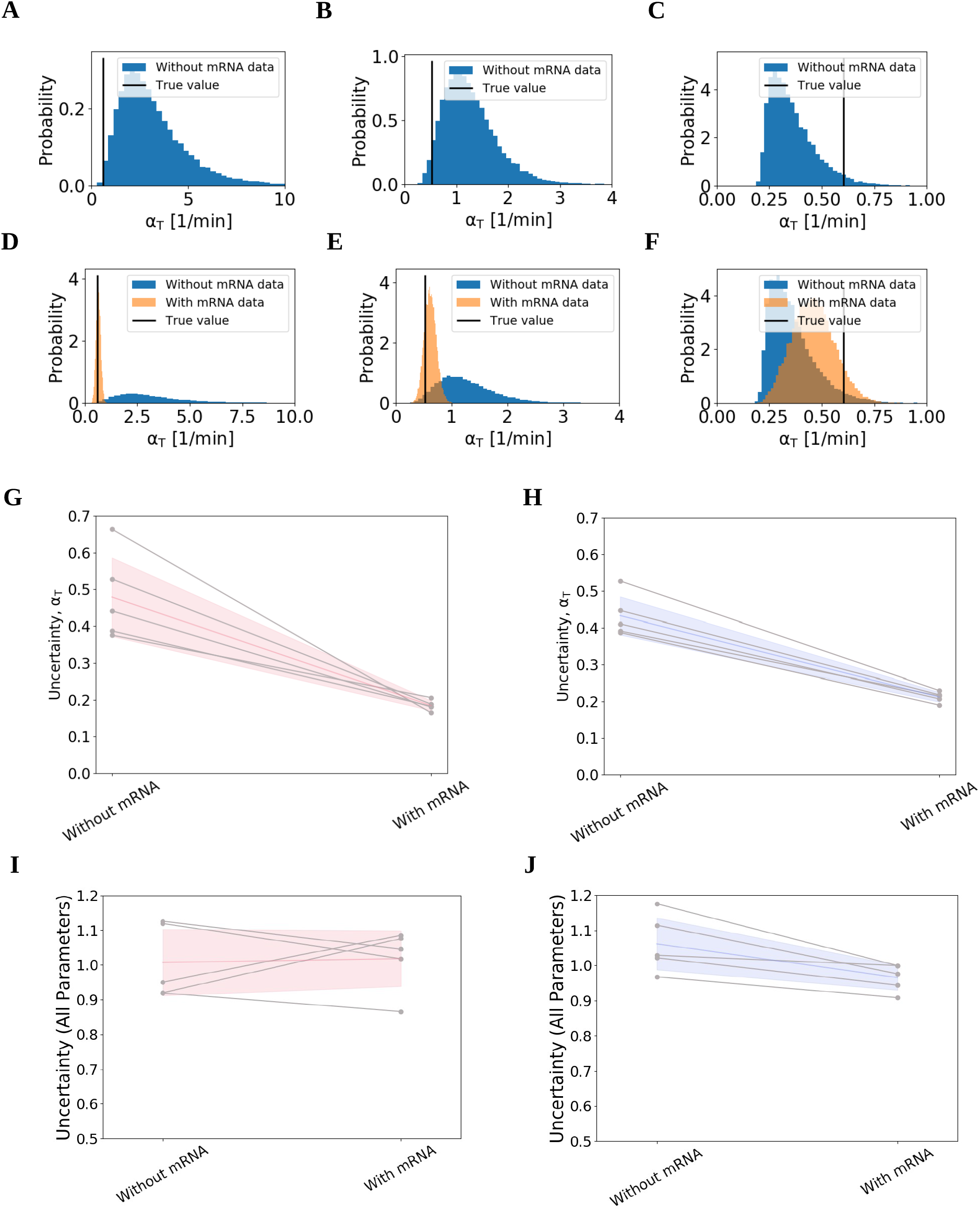
Additional measurements of mRNA copy numbers improve estimates of the average transcription rate. **A,B,C**. Posterior distributions of the average rate of transcription, *α*_*T*_, calculated using the posterior samples of three example data sets from Figure 5. The ground truth value (vertical black line) is poorly estimated by these posteriors. **D,E,F**. The same three posterior distributions for *α*_*T*_ as in A–C, this time comparing posterior samples drawn without mRNA information (blue) and with mRNA information (orange). Here the ground truth value (vertical black line) is much more closely inferred when mRNA information is included. **G**. Uncertainty of *α*_*T*_ for the low coherence data sets from Figure 5. Uncertainty in *α*_*T*_ is reduced by more than 60% when mRNA information is included. Uncertainty on *α*_*T*_ is calculated using the coefficient of variation, defined by the posterior standard deviation over the posterior mean, 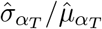 **H**. Same as in G but for high coherence data sets. Here mRNA information reduces uncertainty in *α*_*T*_ by more than 50%. **I**. Values of relative uncertainty RU_***θ***_ (equation (S35) in supplementary Section S.7) for low coherence data sets (cf. Figure 5) with and without additional data on mRNA copy numbers. **J**. RU_***θ***_ for high coherence data sets from Figure 5 with and without additional data on mRNA copy numbers. In G–J the coloured lines and shaded areas represent the mean and standard deviation across observed values, respectively. Data sets in A–C are all sampled for a duration of 12 hours, and with sampling intervals of 8 minutes, 8 minutes, and 5 minutes respectively. All data sets considered in G–J are sampled every 15 minutes for a duration of 12 hours and the parameters correspond to the low and high coherence parameter sets respectively (see Table S2).

Hence we next assume that, in addition to data on the dynamics of protein expression, measurements of mRNA copy numbers have been conducted on the observed cells. Specifically, we generate *in silico* data mimicking a single-molecule *in situ* hybridisation (smFISH) experiment. Such sm-FISH experiments generate distributions of mRNA copy numbers, thus providing a snapshot of mRNA levels across a population at a fixed time point^[66;67]^. To account for this additional data, we incorporate the observed distribution of mRNA copy numbers into our likelihood function, such that it effectively penalises parameters for which inferred copy numbers of mRNA are outside the experimentally observed range (see supplementary Section S.9).

We find that this inclusion of mRNA information collected from a cell population leads to more accurate inference of the average transcription rate for single cells, using our algorithm. Observing example data sets from Figure 5, the posterior distributions cover multiple orders of magnitude if only protein expression data is considered in Figure 6D,E, with the mean of the distribution being 5.4 times larger than the true value in Figure 6D, and 2.4 times larger in Figure 6E, respectively. Upon inclusion of mRNA information, these posterior distributions are instead concentrated around the true value, with a relative error below 15.3%. In both examples the ground truth is contained within the 65% HDI. In Figure 6F a posterior distribution that is already close to the true value gets further constrained by the additional mRNA data. In these examples, the observed reduction of uncertainty on the inferred transcription rate is accompanied by a reduction of uncertainty on estimated mRNA copy numbers for individual cells, as inferred by the Kalman filter (see e.g. supplementary Figure S5A vs. S5B). Investigating the uncertainty on the average inferred transcription rate across data sets introduced in Figure 5, we observe a reduction in uncertainty of 61.8% for low coherence parameter combinations (Figure 6G), and a reduction in uncertainty of 51.2% for high coherence parameter combinations (Figure 6H).

How does this improved estimate of transcription rate affect overall uncertainty across parameter space, as analysed in Figure 5? Counter-intuitively, we find that this inclusion of mRNA data into our parameter inference does not reduce overall parameter uncertainty (Figure 6I,J). For data sets from the low coherence parameter combination, the relative uncertainty increases by 1.1% on average when mRNA information is included (Figure 6I). For data sets from the high coherence parameter combination, uncertainty decreases slightly (9.1% on average (Figure 6J)). Importantly, this reduction of uncertainty is considerably smaller than the reduction of uncertainty observed when longer measurement durations are considered (cf. Figure 5D). We make analogous observations as inference accuracy is analysed (Figure S4), instead of uncertainty. Inference accuracy is not reduced for high coherence data sets when data on mRNA copy numbers is included, and it is only slightly reduced for some of the low coherence data sets, with the effect being much smaller than the effect of considering longer time course data (cf. Figure S3).

The effect that overall uncertainty is not decreased as new data on mRNA copy numbers is included contradicts the intuition that more accurate inference of the average rate of transcription *α*_*T*_ will also reduce uncertainty on model parameters regulating *α*_*T*_, such as the basal transcription rate, *α*_*m*_ and the repression threshold, *P*_0_. This effect may be attributed to correlations between these parameters, which we typically observe in our posterior distributions (see Figure 3F). For the data set in Figure 6A, inference of *α*_*T*_ is improved upon inclusion of the mRNA. This leads to a tighter coupling between the parameters *α*_*m*_ and *P*_0_ (Figure S6). However, this constraining of the posterior distribution is not reflected in either of the marginal posterior distributions. Thus, although the inclusion of *in silico* smFISH data reduces the spread of the posterior distribution overall, uncertainty within the marginal posterior distributions is not reduced, and individual parameter estimates are not improved. An additional factor is that data from smFISH experiments may be considered to reflect the time-averaged mRNA copy number distribution of single cells. Hence, these data might not reduce uncertainty on parameters that are expected to predominantly alter the dynamics rather than the level of expression, such as the transcriptional delay *τ* and Hill coefficient *h*. Hence, to better infer these parameters, other strategies, e.g. those discussed in Figure 5, may be required.

We conclude that distributions of mRNA copy numbers from population-level measurements can be used to infer average transcription rates for individual cells, using our inference method, which may facilitate the study of transcriptional dynamics in the context of gene expression oscillations. Together with results from Figure 5, this illustrates how our method may be used to evaluate the benefit of different experiments *in silico*, and highlights that our method can be naturally extended to utilise additional data of different types.

## 4 Discussion

The aim of this work was to develop a statistical tool that can be used to infer kinetic parameters of the auto-negative feedback motif, based on typically collected protein expression time series data from single cells. Importantly, the stochastic nature of the involved processes demanded a method that enables uncertainty quantification. We have achieved our aim by embedding a non-linear delay-adapted Kalman filter into the MALA sampling algorithm. Our method can generate accurate posterior distributions for the simultaneous inference of multiple parameters of the auto-negative feedback motif. The produced posterior distributions are more informative than those from previous approaches. Since our method can be applied to data from single cells, it enables the investigation of cell-to-cell heterogeneity within cell populations. It can further be used to make experimental design recommendations, which we demonstrated by investigating how parameter uncertainty may depend on the position in parameter space, the sampling frequency, and the length of the collected time series data. Additionally, our method may be extended to account for the presence of different types of data, for example to improve estimates of the transcription rate for individual cells.

Often, new inference algorithms are presented on a single data set, and due to necessary tuning requirements of the involved sampling methods, further data sets are not considered. However, it is important to understand the behaviour of a method for a range of data sets if we wish to make experimental design recommendations. It is an achievement of this paper that we provide a method that demonstratively can reliably infer parameters, even when the size and structure of the data are changed significantly.

The mathematical model underlying our method aims to describe the dynamic expression of a protein which is controlled by auto-negative feedback. The success of our inference relies upon how well this model approximates reality. Mathematical models for the oscillatory expression of transcription factors are informed by experimental research^[57;68]^ and have been developed over time^[3;8;22;23;64]^. Existing model extensions include interactions with other genes or microRNAs^[21]^ and future models could include effects of transcriptional bursting^[69]^. The simple model used here provides a starting point for developing inference algorithms for further models including non-linear, stochastic interactions as well as delays, and future validation of experimental predictions can be used to guide data-driven model improvements. To this end, our algorithm may enable model selection to identify gene regulatory network properties, such as interactions between multiple transcription factors.

Chemical Langevin Equations such as eqs. (2) and (3) approximate the full stochastic dynamics of the system by assuming Gaussian increments. Further, our Kalman filter assumes that measurement errors follow a Gaussian distribution, and are not correlated between consecutive time points. The likelihood calculations within the Kalman filter assume that distributions of protein copy numbers, which are predicted by eqs. (2) and (3), can be approximated by Gaussian distributions.

The Gaussian approximation within the Chemical Langevin Equation can break down when molecule concentrations are very low, resulting in an inaccurate simulation of the dynamics. We do not expect this to be a problem for data analysed in this paper, since protein copy numbers throughout our analysis are around 50,000 protein molecules per nucleus. In other applications, the validity of the Chemical Langevin Equation may be explicitly tested on samples from the posterior distribution by directly comparing simulated expression time series with those obtained from an exact sampling algorithm, such as the Gillespie algorithm^[70]^. Similarly, simulations of the Chemical Langevin Equation can be used to test assumptions on the Gaussianity of the state space made within the Kalman filter. In cases where these assumptions do not hold, alternative inference algorithms, such as particle filter methods, may need to be developed.

For Bayesian inference problems it is common to use MCMC samplers, such as MH or MALA. We have found that combining a delay-adapted non-linear Kalman filter and MALA can allow us to infer parameters of the auto-negative feedback motif. This builds on previous approaches which applied a Kalman filter in the context of a different transcriptional feedback motif with delay^[52]^. MCMC algorithms typically require tuning which may be data specific. We have taken steps to reduce additional input from the user by using MALA, which proposes new samples based on the gradient of the target posterior, hence accounting for geometric properties of parameter space, which can result in faster, more robust performance on some distributions^[44]^. MALA also has fewer tuning parameters than other algorithms, such as HMC. This allows us to more easily incorporate adaptation into our algorithm^[71]^. Surprisingly, the MALA sampler did not result in faster convergence than MH on example posteriors from our model (see Supplementary Figure S7). Hence, the added computational cost of calculating likelihood gradients will not be beneficial in all applications, especially since, in our model, gradient calculations increase the computational cost of individual parameter samples by a factor 12. We expect the availability of likelihood gradients to achieve a speed-up in high-dimensional problems, where convergence speeds of MALA scale with *d*^1*/*3^, rather than *d*^1^ for MH^[72]^, for model dimension *d*. Note, that more efficient MCMC algorithms can eliminate the problem of tuning entirely^[44]^. These methods rely on the computation of the Hessian, i.e. the second derivative of the likelihood function. Deriving expressions for the Hessian and investigating the efficiency of the resulting algorithm is thus a potential avenue for future work. In our applications of the algorithm to experimentally measured data, we detrended the data before applying our inference (Figure 4B). Such detrending is commonly used when analysing time series of oscillatory signals^[2;73;74]^. The detrending removes signal fluctuations from the recorded time series that vary on a much longer time scale than the ultradian oscillations that are being considered. This is necessary, since our model cannot describe such long-term fluctuations. Specifically, independently of the model parameter chosen, simulated traces from the Chemical Langevin Equation (eqs. (2) and (3)) do not include long-term trends. Hence, detrending prevents any bias that the presence of a long-term trend in the data may introduce to the parameter inference. When the algorithm will be applied to data from other transcription factors, we recommend to exclude data that contains trends with timescales that are longer than the fluctuations and oscillations that are expected to emerge from the auto-negative feedback, in line with previous detrending recommendations^[73;74]^. Presumably, variations in the long-term trend of the data stem from a time-dependence of one or multiple of the model parameters due to regulatory processes that our model does not account for. Hence, future improvements to our algorithm may be developed where the temporal variation of model parameters is inferred, instead of one static value.

When applying our inference method to experimental data (Figure 4), we relied on previously reported values for the measurement variance, Σ_*ϵ*_, in the data set that we considered^[2]^. When users seek to apply our algorithm to other data where previously published values are not available for Σ_*ϵ*_, this parameter can be inferred following the same procedure as reported in Manning et al. (2019).

Our algorithm opens up the investigation of research problems, such as cell-to-cell heterogeneity in dynamic gene expression, which would previously not have been accessible. In future applications, our algorithm may provide a non-invasive method to measure the kinetic parameters of the gene of interest, such as the translation and transcription rates, or properties of the gene’s promoter, which are described by the repression threshold and Hill coefficient parameters in our model. On experimental data sets where multiple, qualitatively different dynamics are observed^[75–77]^, our algorithm may provide insights into the mechanistic origin of these different dynamics, by identifying differences in inferred parameter values between the observed cells or cell populations. In order to classify whether observed differences between posterior distributions are significant, one can construct the posterior distribution describing the difference between parameter values from both cells or populations, and test whether the credible interval of that distribution contains zero^[78]^. To facilitate such analysis, our method may for example be combined with clustering algorithms on the time series data, such as Gaussian mixture modelling. Since different dynamic patterns of gene expression have been observed in multiple studies of auto-repressing transcription factors^[2;3]^, we anticipate that these approaches will spark new scientific investigations.

Throughout, we have assumed that measurements in the form of protein copy numbers per nucleus are available over time. To collect such data, it is necessary to combine endogenous fluorescent reporters with FCS in order to translate reporter intensity values to molecule concentrations. Future versions of our algorithm may be applicable to data where FCS is not available, if one extends our measurement model (*F*, Section 2.2) to include an unknown, linear scaling parameter between protein copy numbers and imaged intensity values.

We highlight that the impact of this work is not limited to a single gene in a single model system. The conceptual framework and derivations described here are applicable to any system which can be described by delayed stochastic differential equations, although there may be computational limitations as model sizes increase.

## Supporting information

Supplementary methods and seven supplementary figures

## Ethics statement

Animal experimentation: All animal work was performed under regulations set out by the UK Home Office Legislation under the 1986 United Kingdom Animal Scientific Procedures Act.

## Data accessibility

All code and data used in this article are freely available on our github repository, https://github.com/kursawe/hesdynamics.

## Authors’ contributions

J.B., C.M., M.R., N.P. and J.K. conceived the study and contributed to the manuscript. J.B. developed and implemented the algorithm, ran the simulations, performed the analysis, and produced the figures. J.K. contributed to the algorithm development. M.R., N.P. and J.K. supervised the study. J.B. and J.K. co-wrote the manuscript with input from C.M., M.R. and N.P.. C.M. provided microscopy data and FCS analysis. All authors gave final approval for publication and agree to be held accountable for the work performed therein.

## Acknowledgements

The authors would like to acknowledge the use of the Computational Shared Facility at The University of Manchester.

