## Supplementary methods and seven supplementary figures for "Inferring kinetic parameters of oscillatory gene regulation from single cell time series data"

Here we provide full derivations for the prediction and update step in the non-linear, delay-adapted Kalman filter. We detail the MCMC sampling algorithms discussed in the main text, as well as our chosen method for adaptive step size selection within the MCMC. Then, we show how the gradient of the likelihood can be computed within the Kalman filter. We further describe our process for choosing prior distributions, and for assessing convergence of our Markov chains, before discussing how oscillation quality of time series data is quantified, and how the likelihood function is extended to account for data on mRNA copy numbers. Finally, we give details about the data collection and analysis. Supplementary figures are provided at the end.

### S.1 Kalman filter prediction and update step

In order to predict the probability distribution over the state space $\mathbf{X}$ at observation $k$, it is necessary to calculate $\rho_t$ and $P_t$ as defined in the main manuscript (Section 2.2). Here, we show how we derive delay differential equations for both quantities, closely following Calderazzo et al. (2018).

**Prediction Step** We first rewrite eqs. (2) and (3) in the general form of a vector-valued, delayed stochastic differential equation:

$$\begin{aligned} d\mathbf{X}(t) &= g(\mathbf{X}(t))dt + f\left(\mathbf{X}(t-\tau)\right)dt \\ &\quad + \sqrt{l(\mathbf{X}(t)) + q\left(\mathbf{X}(t-\tau)\right)}dB(t), \end{aligned} \tag{S10}$$

where $B(t)$ is a 2-dimensional Wiener process, and

$$g(\mathbf{X}(t)) = \begin{bmatrix} -\mu_m m(t) \\ \alpha_p m(t) - \mu_p p(t) \end{bmatrix},$$

$$f(\mathbf{X}(t-\tau)) = \begin{bmatrix} \alpha_m f(p(t-\tau)) \\ 0 \end{bmatrix},$$

$$l(\mathbf{X}(t)) = \begin{bmatrix} \mu_m m(t) & 0 \\ 0 & \alpha_p m(t) + \mu_p p(t) \end{bmatrix},$$

$$q(\mathbf{X}(t-\tau)) = \begin{bmatrix} \alpha_m f(p(t-\tau)) & 0 \\ 0 & 0 \end{bmatrix}.$$

Next, we aim to use this definition to derive delay differential equations for $\rho$ and $P$. To do so, we employ Taylor expansions. The first order Taylor expansions of $g(\cdot)$ and $f(\cdot)$ about $\rho_t$ are

$$g(X_t) \approx g(\rho_t) + J_g(\rho_t) \cdot (X_t - \rho_t),$$

and

$$f(X_{t-\tau}) \approx f(\rho_{t-\tau}) + J_f(\rho_{t-\tau}) \cdot (X_{t-\tau} - \rho_{t-\tau}).$$

We can hence write

$$X_{t+\delta_t} = X_t + g(X_t)\delta_t + f(X_{t-\tau})\delta_t + \sqrt{l(X_t) + q(X_{t-\tau})}\mathcal{N}(0, \delta_t) + \mathcal{O}(\delta_t^2), \tag{S11}$$

where $\mathcal{N}(0, \delta_t)$ is a Gaussian distribution with mean zero and variance $\delta_t$. Inserting this into the definition of $\rho_t$, we find

$$d\rho_t = g(\rho_t)dt + f(\rho_{t-\tau})dt. \tag{S12}$$

Similarly, we insert eq. (S11) into the definition of $P$ to find

$$\begin{aligned} dP_t = & \left[J_g(\rho_t)P_t + P_t^T J_g(\rho_t)^T\right] dt \\ & + \left[J_f(\rho_{t-\tau})P_{t-\tau,t}\right] dt \\ & + \left[P_{t,t-\tau} J_f(\rho_{t-\tau})^T\right] dt \\ & + A\left(\rho_t, \rho_{t-\tau}\right) dt, \end{aligned} \tag{S13}$$

where $A(\rho_t, \rho_{t-\tau}) = l(\rho_t) + q(\rho_{t-\tau})$ and $P_{t,s} = \mathrm{Cov}(\mathbf{X}(t), \mathbf{X}(s) \mid y_{0:t-1})$.

Lastly, we have

$$dP_{s,t} = P_{s,t}J_g^T(\rho_t)dt + P_{s,t-\tau}J_f^T(\rho_{t-\tau})dt. \tag{S14}$$

We can then use a standard forward Euler method with a time step which represents the absolute time difference between consecutive hidden states, to numerically integrate these delay differential equations between observations. Before the first prediction step can be executed, it is necessary to define initial and past conditions for these delay differential equations. The state space mean is initialised at negative times with the steady state solutions to the deterministic approximation of the model, i.e. eqs (2) and (3) of the main manuscript without their respective noise terms. At steady state, we have

$$p^* = \frac{\alpha_m \alpha_p}{\mu_m \mu_p} f(p^*) \tag{S15}$$

and

$$m^* = \frac{\mu_p}{\alpha_p} p^*. \tag{S16}$$

The state space variance is initialised with a multiple of this mean. Specifically, the variance of the mRNA is initialised at 20 times its mean value, and the variance of the protein is initialised at 100 times its mean value. These are chosen to reflect the variance of protein and mRNA expression from the Chemical Langevin Equations. For simplicity, off-diagonal elements of the state space covariance are initialised at zero. We expect the influence of these initial and past conditions to diminish after the first update step has been applied and as the number of data points increases.

**Update Step** For our update step we derive an expression for the mean and variance of the state space distribution $\pi(x_{t-\tau:t} \mid y_{0:t})$, denoted $\rho^*_{t-\tau:t}$ and $P^*_{t-\tau:t}$ respectively. That is, the likelihood of our state space estimates from the past time $t-\tau$ to the current time, $t$, given all of our current observations. This is necessary in order to accurately predict the state space distribution at the next observation time point, $\pi(x_{t+\Delta t} \mid y_{0:t})$, as past states can affect future states due to the presence of delays.

Assuming approximate Gaussianity of the joint distribution $\pi(x_{t+\Delta t}, y_{t+\Delta t} \mid y_{0:t})$, following Särkkä (2013), we can obtain the following expression for the updated state space mean and variance:

$$\rho^*_{t-\tau:t} = \rho_{t-\tau:t} + K_{t-\tau:t}\left(y_t - F\rho_t\right), \tag{S17}$$

$$P^*_{t-\tau:t} = P_{t-\tau:t} - K_{t-\tau:t} S_t K^T_{t-\tau:t}, \tag{S18}$$

$$K_{t-\tau:t} = P_{t-\tau:t,t} F^T S_t^{-1}, \tag{S19}$$

where $S_t = F P_t F^T + \Sigma_\epsilon$.

### S.2 Markov Chain Monte Carlo sampling algorithms

For completeness, we provide here the algorithms for the Metropolis-Hasting random walk, and the Metropolis-adjusted Langevin Algorithm.

#### Metropolis-Hastings Random Walk

We initialise the algorithm with a set of the model parameters, $\boldsymbol{\theta}_0$, chosen by minimising the log likelihood function using Powell's method[3]. In the case of our model, each parameter vector is composed as $\boldsymbol{\theta} = [P_0, h, \mu_m, \mu_p, \alpha_m, \alpha_p, \tau]^T$. At each iteration, $i$, we propose a combination of parameters, $\boldsymbol{\theta}'$, from a symmetric distribution, $q(\cdot)$, centered around the current parameters, $\boldsymbol{\theta}_i$. In this paper we use the normal distribution $\mathcal{N}(\boldsymbol{\theta}, \epsilon\Sigma)$, where $\epsilon$ is called the step size, and $\Sigma$ is the proposal covariance matrix. An acceptance ratio, $\alpha(\boldsymbol{\theta}_i, \boldsymbol{\theta}')$, defined by

$$\begin{aligned}\alpha(\boldsymbol{\theta}_i, \boldsymbol{\theta}') &= \min\left\{1, \frac{\pi(\boldsymbol{\theta}' \mid \mathbf{y})}{\pi(\boldsymbol{\theta}_m \mid \mathbf{y})}\right\} \\ &= \min\left\{1, \frac{\pi(\mathbf{y} \mid \boldsymbol{\theta}')\pi(\boldsymbol{\theta}')}{\pi(\mathbf{y} \mid \boldsymbol{\theta}_i)\pi(\boldsymbol{\theta}_m)}\right\}\end{aligned}$$

is calculated, and a uniform random number, $u$, is generated on $[0, 1]$.

1. If $u \leq \alpha(\boldsymbol{\theta}_i, \boldsymbol{\theta}')$, we *accept* the proposal $\boldsymbol{\theta}'$ and set $\boldsymbol{\theta}' = \boldsymbol{\theta}_{i+1}$.

2. If $u \geq \alpha(\boldsymbol{\theta}_i, \boldsymbol{\theta}')$, we *reject* the proposal $\boldsymbol{\theta}'$ and set $\boldsymbol{\theta}_t = \boldsymbol{\theta}_{i+1}$.

The distribution of random draws $\{\boldsymbol{\theta}_i \mid i \in \{0, 1, \ldots, N_s\}\}$ converges to our posterior distribution, $\pi(\boldsymbol{\theta} \mid \mathbf{y})$, as $N_s \to \infty$. In practice, the number of iterations necessary for accurate inference depends on a number of factors, including the complexity of the problem, the number of parameters to be inferred, and the relationship between the posterior and proposal distributions. We outline our choice of $N_s$ in Section S.3

#### Metropolis-adjusted Langevin Algorithm (MALA)

Whilst the MH algorithm is relatively simple to understand and implement, it can fail to identify the posterior if it has a non-trivial, curved correlation structure[4], or if it is multi-modal[5]. It can also become slow when dimensionality is high, i.e. when there are a large number of parameters to infer. In these cases the algorithm typically requires a very large number of samples, which is time consuming, especially if the likelihood calculation is slow. One way to address this is to use a sampling algorithm that uses the gradient of the posterior distribution, as well as it's absolute values, to draw new samples. The gradient can help guide the search towards the modes of the posterior distribution, making for a more efficient algorithm which typically needs fewer samples. A number of different MCMC approaches utilise the gradient of the distribution, such as MALA, and the NO-U-Turn sampler [6], which is an extension on the original Hamiltonian Monte Carlo (HMC) [7].

We use MALA, since it is well-established that it has better convergence properties than the MH algorithm [8]. Our algorithm for MALA iterates over the same steps as the MH algorithm, with an adjusted expression for the proposed parameter combination. Where MH uses the proposal

$$\boldsymbol{\theta}' = \boldsymbol{\theta} + \sqrt{\epsilon}\Sigma^{1/2}\xi, \tag{S20}$$

MALA instead uses

$$\boldsymbol{\theta}' = \boldsymbol{\theta} - (\epsilon/2)\Sigma U(\boldsymbol{\theta}) + \sqrt{\epsilon}\Sigma^{1/2}\xi, \tag{S21}$$

where $\xi \sim \phi(0, I_d)$, with $I_d$ denoting the $d$-dimensional identity matrix, $d$ is the dimension of the posterior distribution, and $U(\boldsymbol{\theta}) = -\nabla \log \pi(\mathbf{y} \mid \boldsymbol{\theta})$.

### S.3 Implementation of adaptive sampling strategies

We seek to generate $N_s$ independent samples that accurately characterise our posterior distribution, but samples are dependent on the starting point of the Markov chain. As increasingly many samples are generated, this dependence diminishes. The correlation between samples can be reduced if an appropriate covariance matrix is chosen for the proposal distribution. One suitable covariance matrix is the covariance of the target posterior distribution, which may be estimated from samples of the MCMC itself. To do so, we *warm up* the Markov chains in two steps by drawing $N_{w_1}$ samples and discarding the first $N_{b_1}$ samples as *burn-in*. With the remaining samples we construct a proposal covariance matrix, and repeat the process with $N_{w_2}$ samples and a burn-in of size $N_{b_2}$. The adaptive MCMC is then started using the covariance matrix estimated in this way as the proposal covariance matrix. We adaptively update our step size at iteration $i$, $\epsilon_i$, using a modified version of Algorithm

4 from Andrieu and Thoms (2008).

At each iteration $i$, the step size is updated using

$$\gamma_1 = \frac{1}{i^{c_1}},$$
$$\gamma_2 = c_0\gamma_1,$$
$$\log(\epsilon_{i+1}^2) = \log(\epsilon_i^2) + \gamma_2(\alpha_i - 0.574), \tag{S22}$$

where $c_0 = 1$, $c_1 = \log(10)/\log(N_s/5)$, and $\alpha_i$ is the current acceptance probability. We use 0.574 as the target acceptance rate for MALA, as shown above, and 0.234 for MH[10].

In our experience, this approach results in far better performance than without warm up or adaptively updating the step size. In practice, we found that consistent posteriors are obtained by running 8 Markov chains in parallel, where each chain has $N_s = 80,000$, $N_{w_1} = 0.3 \times N_s$, $N_{w_2} = 0.7 \times N_s$, and $N_{b_i} = 0.5 \times N_{w_i}$, $i = 1, 2$. Warm up samples from all eight chains are pooled to calculate the covariance matrix for the final run of MCMC. This approach to warm up is used in the same way when either sampler, MH or MALA, is chosen.

### S.4 Computing the gradient of the log-posterior

The Kalman filter has previously been used within gradient based sampling schemes[11], which illustrated that it is possible to calculate the gradient with an iterative scheme similar to the scheme for the direct likelihood calculation. In the following, we show that these previous approaches can be extended to derive the gradient of the delay-mediated Kalman filter used here.

For MALA, we compute the derivative of the negative log-posterior $U = -\log(\pi(\boldsymbol{\theta}|\mathbf{y})$ with respect to the parameters, $\boldsymbol{\theta}$,

$$\begin{aligned}\frac{\partial U(\boldsymbol{\theta})}{\partial \boldsymbol{\theta}} &= \frac{\partial(-\log(\pi(\mathbf{y} \mid \boldsymbol{\theta})\pi(\boldsymbol{\theta})))}{\partial \boldsymbol{\theta}} \\ &= \frac{\partial\left(-\log\left(\prod_{t\in\mathcal{T}} \phi(y_t; F\rho_t, FP_tF^T + \Sigma_\epsilon)\pi(\boldsymbol{\theta})\right)\right)}{\partial \boldsymbol{\theta}}\end{aligned}$$

$$= -\frac{\partial}{\partial \boldsymbol{\theta}} \log \left( \prod_{t \in \mathcal{T}} \frac{\exp\left(-\frac{1}{2}(y_t - F\rho_t)^T (FP_tF^T + \Sigma_\epsilon)^{-1}(y_t - F\rho_t)\right)\pi(\boldsymbol{\theta})}{\sqrt{\det(2\pi(FP_tF^T + \Sigma_\epsilon))}} \right)$$

$$= \frac{\partial}{\partial \boldsymbol{\theta}} \left[ \sum_{t \in \mathcal{T}} \frac{1}{2} \log\left(\det(2\pi S_t(\boldsymbol{\theta}))\right) + \right.$$

$$\left. \frac{1}{2} \sum_{t \in \mathcal{T}} (y_t - F\rho_t(\boldsymbol{\theta}))^T (S_t(\boldsymbol{\theta}))^{-1} (y_t - F\rho_t(\boldsymbol{\theta})) - \log(\pi(\boldsymbol{\theta})) \right],$$

where $S = FP_tF^T + \Sigma_\epsilon$ has been used in the last line, and the dependence of $\rho$ and $S$ on $\boldsymbol{\theta}$ has been emphasized. $\mathcal{T}$ denotes the set of time points for each observation, $\{i \cdot l \mid i \in \{0, 1, \ldots, n-1\}\}$. For a single parameter this simplifies to

$$\begin{aligned} \frac{\partial U(\boldsymbol{\theta})}{\partial \theta_k} &= \frac{1}{2} \sum_{t \in \mathcal{T}} \left( \mathrm{Tr}\left( S_t^{-1} \frac{\partial S_t}{\partial \theta_k} \right) \right) - \frac{1}{2} \sum_{t \in \mathcal{T}} \left( F \frac{\partial \rho_t}{\partial \theta_k} \right)^T S_t^{-1} (y_t - F\rho_t) \\ &- \frac{1}{2} \sum_{t \in \mathcal{T}} (y_t - F\rho_t)^T S_t^{-1} \frac{\partial S_t}{\partial \theta_k} S_t^{-1} (y_t - F\rho_t) \\ &- \frac{1}{2} \sum_{t \in \mathcal{T}} (y_t - F\rho_t)^T S_t^{-1} F \frac{\partial \rho_t}{\partial \theta_k} - \frac{\partial(\log(\pi(\boldsymbol{\theta})))}{\partial \theta_k}. \end{aligned} \tag{S23}$$

This expression can be explicitly calculated if $\mathrm{d}\rho_t/\mathrm{d}\theta_k$ and $\mathrm{d}P_t/\mathrm{d}\theta_k$ are known at each observation time point $t$. These quantities are the predicted state space mean and variance in the prediction step of the Kalman filter. This implies that eq. (S23) can be calculated if the prediction step is adjusted to predict the derivative of $\rho$ and $P$ in addition to their absolute values. To enable this adjustment of the prediction step, we derive differential equations for $\frac{\mathrm{d}}{\mathrm{d}t}(\mathrm{d}\rho_t/\mathrm{d}\theta_k)$ and $\frac{\mathrm{d}}{\mathrm{d}t}(\mathrm{d}P_t/\mathrm{d}\theta_k)$:

$$\begin{aligned} \frac{\mathrm{d}}{\mathrm{d}t} \left( \frac{\partial \rho_t}{\partial \theta_k} \right) &= \frac{\partial}{\partial \theta_k} \left( \frac{\mathrm{d}\rho_t}{\mathrm{d}t} \right) \\ &= \frac{\partial}{\partial \theta_k} \left( g(\rho_t) + f(\rho_{t-\tau}) \right) \\ &= \frac{\partial g(\rho_t)}{\partial \rho_t} \frac{\partial \rho_t}{\partial \theta_k} + \frac{\partial g(\rho_t)}{\partial \theta_k} + \frac{\partial f(\rho_{t-\tau})}{\partial \rho_{t-\tau}} \frac{\partial \rho_{t-\tau}}{\partial \theta_k} + \frac{\partial f(\rho_{t-\tau})}{\partial \theta_k} \\ &= J_g(\rho_t) \frac{\partial \rho_t}{\partial \theta_k} + \frac{\partial g(\rho_t)}{\partial \theta_k} + J_f(\rho_{t-\tau}) \frac{\partial \rho_{t-\tau}}{\partial \theta_k} + \frac{\partial f(\rho_{t-\tau})}{\partial \theta_k}, \end{aligned} \tag{S24}$$

and

$$
\begin{aligned}
\frac{\mathrm{d}}{\mathrm{d}t}\left(\frac{\partial P_t}{\partial \theta_k}\right) &= \frac{\partial}{\partial \theta_k}\left(\frac{\mathrm{d}P_t}{\mathrm{d}t}\right) \\
&= \frac{\partial}{\partial \theta_k}\Bigg(J_g(\rho_t)P_t + P_t^T J_g(\rho_t)^T \\
&\qquad + J_f(\rho_{t-\tau})P_{t-\tau,t} + P_{t,t-\tau}J_f(\rho_{t-\tau})^T \\
&\qquad + l(\rho_t) + q(\rho_{t-\tau})\Bigg) \\
&= J_g(\rho_t)\frac{\partial P_t}{\partial \theta_k} + \frac{\partial P_t^T}{\partial \theta_k}J_g(\rho_t)^T + J_f(\rho_{t-\tau})\frac{\partial P_{t-\tau,t}}{\partial \theta_k} + \frac{\partial P_{t,t-\tau}}{\partial \theta_k}J_f(\rho_{t-\tau})^T \\
&+ \left(\sum_{j=1}^{2}\left(\frac{\partial J_g(\rho_{t_j})}{\partial \rho_{t_j}}\frac{\partial \rho_{t_j}}{\partial \theta_k}\right) + \frac{\partial J_g(\rho_t)}{\partial \theta_k}\right)P_t + P_t^T\left(\sum_{j=1}^{2}\left(\frac{\partial J_g(\rho_{t_j})}{\partial \rho_{t_j}}\frac{\partial \rho_{t_j}}{\partial \theta_k}\right) + \frac{\partial J_g(\rho_t)}{\partial \theta_k}\right)^T \\
&+ \left(\sum_{j=1}^{2}\left(\frac{\partial J_f(\rho_{t-\tau_j})}{\partial \rho_{t-\tau_j}}\frac{\partial \rho_{t-\tau_j}}{\partial \theta_k}\right) + \frac{\partial J_f(\rho_{t-\tau})}{\partial \theta_k}\right)P_{t-\tau,t} \\
&+ P_{t,t-\tau}\left(\sum_{j=1}^{2}\left(\frac{\partial J_f(\rho_{t-\tau_j})}{\partial \rho_{t-\tau_j}}\frac{\partial \rho_{t-\tau_j}}{\partial \theta_k}\right) + \frac{\partial J_f(\rho_{t-\tau})}{\partial \theta_k}\right)^T \\
&+ \left(\sum_{j=1}^{2}\left(\frac{\partial l(\rho_{t_j})}{\partial \rho_{t_j}}\frac{\partial \rho_{t_j}}{\partial \theta_k}\right) + \frac{\partial l(\rho_t)}{\partial \theta_k}\right) + \left(\sum_{j=1}^{2}\left(\frac{\partial q(\rho_{t_j})}{\partial \rho_{t_j}}\frac{\partial \rho_{t_j}}{\partial \theta_k}\right) + \frac{\partial q(\rho_t)}{\partial \theta_k}\right) \qquad \text{(S25)}
\end{aligned}
$$

where $\rho_{t_j}$ means the $j$th element of $\rho_t$. Similarly,

$$
\begin{aligned}
\frac{\mathrm{d}}{\mathrm{d}t}\left(\frac{\partial P_{s,t}}{\partial \theta_k}\right) &= \frac{\partial}{\partial \theta_k}\left(\frac{\mathrm{d}P_{s,t}}{\mathrm{d}t}\right) \\
&= \frac{\partial}{\partial \theta_k}\left(P_{s,t}J_g^T(\rho_t) + P_{s,t-\tau}J_f^T(\rho_{t-\tau})\right) \\
&= \frac{\partial P_{s,t}}{\partial \theta_k}J_g^T(\rho_t) + \frac{\partial P_{s,t-\tau}}{\partial \theta_k}J_f^T(\rho_{t-\tau}) \\
&+ P_{s,t}\left(\sum_{j=1}^{2}\left(\frac{\partial J_g(\rho_{t_j})}{\partial \rho_{t_j}}\frac{\partial \rho_{t_j}}{\partial \theta_k}\right) + \frac{\partial J_g(\rho_t)}{\partial \theta_k}\right)^T
\end{aligned}
$$

$$+ P_{s,t-\tau} \left( \sum_{j=1}^{2} \left( \frac{\partial J_f(\rho_{t-\tau_j})}{\partial \rho_{t-\tau_j}} \frac{\partial \rho_{t-\tau_j}}{\partial \theta_k} \right) + \frac{\partial J_f(\rho_{t-\tau})}{\partial \theta_k} \right)^T . \tag{S26}$$

Similar to the delay differential equations for the state space mean and variance, it is necessary to define values for non-positive times for the delay differential equations provided here before the first prediction step can be executed. To do so, we need to provide derivatives of the initialisation of the state space mean and variance provided in Section S.1. We calculate the derivatives of $p^*$ and $m^*$ with respect to each parameter, $\theta_k$, as follows:

$$\frac{\partial p^*}{\partial \theta_k} = \frac{-\frac{\partial}{\partial \theta_k}\left(\frac{\alpha_m \alpha_p}{\mu_m \mu_p}\right) f(p^*) - \left(\frac{\alpha_m \alpha_p}{\mu_m \mu_p}\right) \frac{\partial}{\partial \theta_k}\left(f(p^*)\right)}{\left(\frac{\alpha_m \alpha_p}{\mu_m \mu_p}\right) \frac{\partial}{\partial p^*} f(p^*) - 1}, \tag{S27}$$

$$\frac{\partial m^*}{\partial \theta_k} = \frac{\partial}{\partial \theta_k}\left(\frac{\mu_p}{\alpha_p} p^*\right). \tag{S28}$$

### S.5 Update step derivatives

The delay differential equations given by eqs. (S24) to (S26) can be used to predict the derivatives of the state space mean and variance with respect to each model parameter, using $\mathrm{d}\rho_t^*/\mathrm{d}\theta_k$ and $\mathrm{d}P_t^*/\mathrm{d}\theta$ as initial (and past) conditions. We incorporate the integration of these ODEs into the prediction step of our Kalman filter. The derivatives of $\mathrm{d}\rho_t^*/\mathrm{d}\theta_k$ and $\mathrm{d}P_t^*/\mathrm{d}\theta$ at each observation can be calculated by extending the update step. Using eqs. (S17) to (S19), we have

$$\begin{aligned}
\frac{\partial \rho^*_{t-\tau:t}}{\partial \theta_k} &= \frac{\partial}{\partial \theta_k}\left(\rho_{t-\tau:t} + K_{t-\tau:t}\left(y_t - F\rho_t\right)\right) \\
&= \frac{\partial \rho_{t-\tau:t}}{\partial \theta_k} + \frac{\partial K_{t-\tau:t}}{\partial \theta_k}\left(y_t - F\rho_t\right) - K_{t-\tau:t} F \frac{\partial \rho_t}{\partial \theta_k}, \quad \text{(S29)} \\
\frac{\partial P^*_{t-\tau:t}}{\partial \theta_k} &= \frac{\partial}{\partial \theta_k}\left(P_{t-\tau:t} - K_{t-\tau:t} S_t K^T_{t-\tau:t}\right) \\
&= \frac{\partial P_{t-\tau:t}}{\partial \theta_k} - \frac{\partial K_{t-\tau:t}}{\partial \theta_k} S_t K^T_{t-\tau:t} - K_{t-\tau:t} \frac{\partial S_t}{\partial \theta_k} K^T_{t-\tau:t} - K_{t-\tau:t} S_t \left(\frac{\partial K_{t-\tau:t}}{\partial \theta_k}\right)^T, \quad \text{(S30)} \\
\frac{\partial K_{t-\tau:t}}{\partial \theta_k} &= \frac{\partial}{\partial \theta_k}\left(P_{t-\tau:t,t} F^T S_t^{-1}\right)
\end{aligned}$$

$$= \frac{\partial P_{t-\tau:t,t}}{\partial \theta_k} F^T S_t^{-1} - P_{t-\tau:t,t} F^T S_t^{-1} \frac{\partial S_t}{\partial \theta_k} S_t^{-1}$$

$$= \frac{\partial P_{t-\tau:t,t}}{\partial \theta_k} F^T S_t^{-1} - P_{t-\tau:t,t} F^T S_t^{-1} F \frac{\partial P_t}{\partial \theta_k} F^T S_t^{-1}. \quad \text{(S31)}$$

Here, $\frac{\partial \rho_{t-\tau:t}}{\partial \theta_k}$ and $\frac{\partial P_{t-\tau:t}}{\partial \theta_k}$ are the derivatives for the state space mean and variance that have been predicted in the previous prediction step. Hence, we can evaluate eq. (S23) for each of the parameters iteratively by evaluating eqs. (S24) to (S26), (S29) and (S30) at each update step, and integrating eqs. (S12) to (S14) within each prediction step of the Kalman filter.

### S.6 Choice of priors and reparameterisation of variables

Bayesian methods require the choice of prior parameter distributions. For the mRNA and protein degradation rates, $\mu_m$ and $\mu_p$, we use fixed values throughout, which are based on previously identified experimental measurements[12]. This facilitates our analysis of experimental time course data from the same paper in Figure 4. For the transcription and translation rates, $\alpha_m$ and $\alpha_p$, we use prior distributions that are log-uniform. For all other parameters, we choose uniform prior distributions. Our chosen parameter ranges are informed by existing literature values[12–17], and summarised in Table S1.

| Parameter | Range of Values |
|---:|:---:|
| Repression threshold, $P_0$ | 0 - 120,000[13] |
| Hill Coefficient, $h$ | 2 - 6[13;14] |
| Protein degradation rate, $\mu_p$ | log(2)/(90 min)[12] |
| mRNA degradation rate, $\mu_m$ | log(2)/(30 min)[12] |
| Basal transcription rate, $\alpha_m$ | 0.01/min - 60/min[15;16] |
| Translation rate, $\alpha_p$ | 1/min - 40/min[17] |
| Transcriptional delay, $\tau$ | 5 min - 40 min[13;14] |

Table S1: The values each of the parameters of the model can take, in relation to the HES5 system, informed by experimental work and biophysical limitations. These values define the prior distributions which we use in our parameter inference algorithms.

The choice of log-uniform priors for the transcription and translation rates, $\alpha_m$ and $\alpha_p$, is motivated by the fact that possible values for these parameters span multiple orders of magnitude, from 0.01 – 120/min and 0.01 – 40/min respectively. To enable sampling from these log-uniform priors, we convert the parameters into logarithmic space using

$$\tilde{\theta} = \ln(\theta),$$

where $\theta$ represents the parameter that is being transformed into logarithmic space, i.e. $\alpha_m$ or $\alpha_p$.

Let $P_1(\theta \mid \mathbf{y})$ and $P_2(\tilde{\theta} \mid \mathbf{y})$ define the posterior distributions for the original and transformed parameter respectively. A transformation of variables changes the shape of the posterior distribution according to

$$P_2(\tilde{\theta} \mid \mathbf{y})\mathrm{d}\tilde{\theta} = P_1(\theta \mid \mathbf{y})\mathrm{d}\theta$$

which in turn gives

$$\begin{aligned} P_2(\tilde{\theta} \mid \mathbf{y}) &= \theta P_1(\theta \mid \mathbf{y}) \\ &= e^{\tilde{\theta}} P_1(e^{\tilde{\theta}} \mid \mathbf{y}), \end{aligned} \tag{S32}$$

since $\dfrac{\mathrm{d}\tilde{\theta}}{\mathrm{d}\theta} = \dfrac{1}{\theta}$.

Using $P_1(\theta \mid \mathbf{y}) = L(\mathbf{y} \mid \theta)\pi_1(\theta)$ and considering a prior that is uniform in logarithmic space, i.e. $\pi_1(\theta) = C/\theta$ with an appropriately chosen constant $C$, eq. (S32) leads to

$$\begin{aligned} \ln(P_2(\tilde{\theta} \mid \mathbf{y})) &= \tilde{\theta} + \ln(L(\mathbf{y} \mid e^{\tilde{\theta}})) + \ln\left(\frac{C}{\theta}\right) \\ &= \ln(L(\mathbf{y} \mid e^{\tilde{\theta}})) + \ln(C) \\ &= \ln(L(\mathbf{y} \mid \theta)) + \ln(C). \end{aligned} \tag{S33}$$

Lastly, taking the derivative gives

$$\frac{\partial}{\partial \tilde{\theta}} \ln(P_2(\tilde{\theta} \mid \mathbf{y})) = \theta \frac{\partial}{\partial \theta} \ln(L(\mathbf{y} \mid \theta)). \tag{S34}$$

### S.7 MCMC success is qualified by multiple convergence diagnostics

When using MCMC samplers such as MH and MALA, it is necessary to check for convergence of the sampled distribution and to identify whether a sufficient number of samples has been generated. A combination of convergence diagnostics give confidence that the obtained samples are useful and represent the posterior accurately. Specifically, we use the split-$\hat{R}$ and effective sample size (ESS) methods[18;19]. We follow the guidance of Vehtari et. al.[19], running multiple chains ($m > 4$), and conclude convergence when split-$\hat{R} < 1.01$ and the total ESS is at least $50 \times 2 \times m$.

When testing our algorithm on *in silico* data, we use test diagnostics to determine inference uncertainty and accuracy. Specifically, for a given set of *in silico* data, let $\mu_{\theta_i}$ define the true value of each parameter that generated the data set, and let $\hat{\mu}_{\theta_i}, \hat{\sigma}_{\theta_i}$ define the posterior mean and standard deviation, respectively, for each parameter. Throughout, we discuss the relative uncertainty (RU)

$$\mathrm{RU}_{\boldsymbol{\theta}} = \sum_{\theta_i} \frac{\hat{\sigma}_i}{|\mathrm{supp}\{\pi(\theta_i)\}|}, \tag{S35}$$

where $|\mathrm{supp}\{\pi(\theta_i)\}|$ is the size of the support of the prior for parameter $\theta_i$. The relative uncertainty hence measures how the marginal standard deviations of the posterior distribution compares to their respective prior widths. Similarly, we introduce the mean error (ME) as

$$\mathrm{ME}_{\boldsymbol{\theta}} = \sum_{\theta_i} \frac{|\mu_{\theta_i} - \hat{\mu}_{\theta_i}|}{|\mathrm{supp}\{\pi(\theta_i)\}|}, \tag{S36}$$

which measures how strongly the inferred posterior mean differs from the ground truth. For any parameters that are sampled in logarithmic space (supplementary Section S.6 above), we consider the samples and prior distributions in the original, *non*-transformed, space.

### S.8 Oscillation quality is quantified by the coherence measure

In the presence of noise, time series can exhibit oscillations of varying quality. Here we introduce a measure to quantify the oscillatory properties of a time series, the coherence. We define coherence as the area under the power spectrum, $f(\omega)$, within a 20% band around the peak frequency, divided by the total area under the curve, following previous approaches[14;20]. The power spectrum is defined by

$$f(\omega) = \langle \hat{p}(w)\hat{p}^*(w) \rangle,$$

where $\hat{p}(\omega)$ is the Fourier transform of the individual protein expression time course data, and $^*$ denotes complex conjugation. Values of the coherence around zero suggest a lack of any periodic behaviour, and values around one suggest sine-wave-like behaviour. For *in silico* data we estimate coherence by averaging the power spectra from 200 traces, which individually were simulated for 8500 minutes. The initial 1000 minutes of the simulation were discarded, and not used in the averaging, to minimise any influence from the initial conditions. The use of the power spectrum to define coherence ensures that coherence is robust to changes in the sampling interval, provided that this interval is shorter than the dominant period of oscillation.

The parameter combinations that correspond to different coherence values in Figure 5 are listed in supplementary Table S2.

| **Coherence value** | $P_0$ | $h$ | $\alpha_m$ | $\alpha_p$ | $\tau$ |
|---|---|---|---|---|---|
| 0.00599 (low) | 88288.602 | 5.589 | 0.644 | 17.321 | 34.0 |
| 0.200 (high) | 30108.800 | 4.950 | 20.006 | 20.540 | 13.0 |

Table S2: **Parameter combinations for specific coherence values** The first row corresponds to 'low' coherence, and the second to 'high' coherence in Figure 5. To simulate noisy measurements, each synthetic data set is generated using a measurement variance of $\Sigma_\epsilon = 10^6$.

### S.9 Extension of the likelihood function to account for data on mRNA expression levels

Our method generates posterior distributions from single-cell time series data of gene expression dynamics, using the likelihood function introduced in section 2.2. It may be desirable to extend this likelihood function to account for further types of data, if these are available, in order to further constrain the model parameters. Here, we briefly describe how our likelihood function can be extended if, in addition to measurements of protein expression over time, data on the population-level distribution of mRNA copy numbers is available. Such data may for example be collected via single-molecule fluorescence *in situ* hybridisation (smFISH) experiments[21;22].

Let us assume a population-level distribution of mRNA copy numbers has been observed, and that this distribution can be approximated by a Gaussian with mean $\hat{\mu}_m$ and variance $\hat{\sigma}_m^2$. We can use these data to define an extended likelihood function as

$$\pi(\mathbf{y}, \hat{\mu}_m, \hat{\sigma}_m^2 \mid \boldsymbol{\theta}) = \pi(\mathbf{y} \mid \boldsymbol{\theta}) \times \phi(\hat{m}; \hat{\mu}_m, \hat{\sigma}_m^2), \tag{S37}$$

where $\hat{m}$ is the time-average of the mean inferred mRNA copy number from the Kalman filter (i.e. the first entry of $\rho_t$ in section 2.2 averaged over all discretisation time points), $\pi(\mathbf{y} \mid \boldsymbol{\theta})$ is defined in eq. (6), and $\phi(\cdot)$ is a Normal density function, defined in eq. (7).

To generate data from *in silico* smFISH experiments, we generate the distribution of observed mRNA copy numbers numerically by simulating data from the same parameter combination that is used to generate a given *in silico* protein expression time series. Specifically, we simulate 5000 traces of time course data with a duration of 6000 minutes each, using equations (2) and (3). The first 1000 minutes of each time series are discarded to avoid any influence of the initial and past conditions. The mean and standard deviation is then calculated from all remaining simulated mRNA values, considering all simulated time points in all time series.

Our extended likelihood function designed in this way penalises parameter combinations for which the mean inferred mRNA copy numbers are outside the experimentally observed range. Hence, we expect our method to be applicable even if the mean mRNA copy number varies between cells. If the experimental setup is able to precisely measure the mean mRNA copy number for a specific time series of protein expression, our population-level variance $\hat{\sigma}_m^2$ may instead be replaced with the variance of the measured mean, thus enabling a more a more restrictive inference.

### S.10 Data collection and analysis

#### S.10.1 Imaging of Primary NS cells

To generate data in Figure 1A, primary NS cells were isolated from the dissected cortex of E12.5 Venus::HES5 embryos[23] and cultured as previously described[24]. 200,000 cells up to passage 15 were plated on laminin (Sigma, UK) coated 35mm glass-bottom dishes (Greiner-Bio One) and imaged in NS proliferation media at 37°C in 5% $CO_2$ using a Plan Fluor 40x 1.3NA oil objective on an Nikon A1-R inverted confocal microscope. A 25$\mu$m $z$-range was used to ensure cells maintained in focus. Maximum projections in $z$ direction were generated before manual cell tracking using Imaris spot function (Bitplane).

#### S.10.2 Conversion of Venus::HES5 intensity to molecule number

Here, we briefly describe how we translate time series of fluorescent intensity from Manning et al. (2019) into time series of molecule numbers. Throughout our analysis, we use time series data of fluorescent intensity that were published by Manning et al. (2019) and which have been corrected for (i) photobleaching and (ii) for weaker intensity signals at deeper $z$-positions of the imaged nuclei. A quantile-quantile plot was generated between the distribution of tracked Venus::HES5 intensities and the distribution of nuclear Venus::Hes5 concentrations across the tissue. The distribution of tracked Venus::Hes5 intensities was generated from cells in tissues with heterozygous Venus::Hes5 reporter. All recorded single cells at all time points were used. The distribution of nuclear Venus::HES5 concentrations was generated using fluorescent correlation spectroscopy (FCS) on cells in tissue slices from the same domain of the E10.5 spinal cord and with a homozygous Venus::Hes5 reporter. Data from from multiple tissue slices and multiple experiments was used. The required FCS data was also published by Manning et al. (2019). Linear regression on the quantile-quantile plot was used to generate a calibration curve between Venus::HES5 intensity and Venus::HES5 concentration over the middle 90% of the range. The gradient of the line was used as a scaling factor and applied to the intensity values in the Venus::HES5 expression time-series to transform intensity to concentrations.

Finally, the average nuclear volume from Manning et al. (2019) was taken into account to convert concentrations into nuclear molecule numbers.

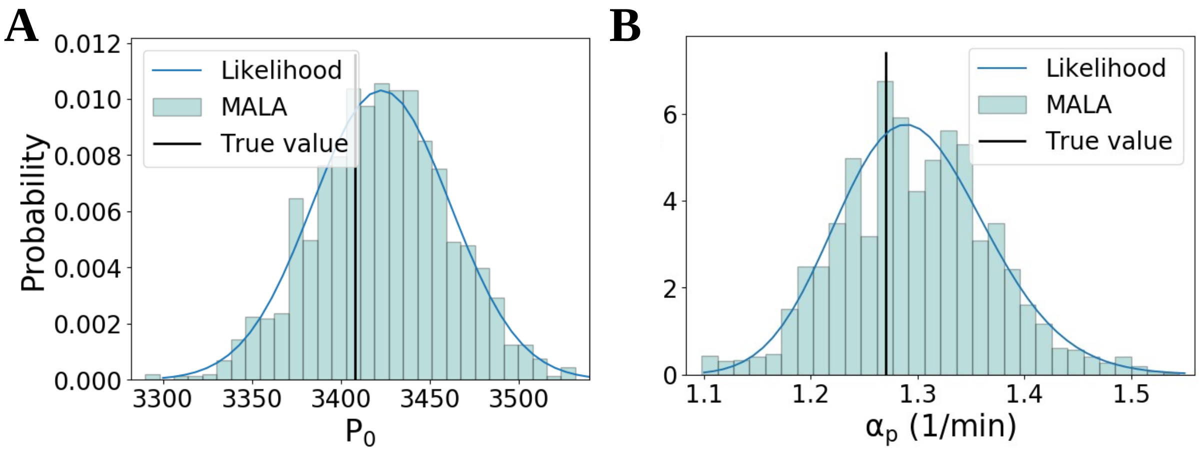


Figure S1: **Our algorithm accurately samples posterior distributions. A, B.** Posterior distributions for one-dimensional inference. For individual model parameters, posterior distributions were inferred while keeping all other parameters fixed, respectively. Shown above are the inferred marginal posteriors for the repression threshold (A) and translation rate (B) respectively as histograms, using MALA as the underlying sampling algorithm for 2500 samples. The blue lines are the analytical likelihood calculations. The sampled and analytical distributions coincide. Other parameters are shown in Figure 2.

| **Parameter** | **True Value** | $\boldsymbol{\mu}$ **(MALA)** | $\boldsymbol{\mu}$ **(MH)** | $\boldsymbol{\sigma}$ **(MALA)** | $\boldsymbol{\sigma}$ **(MH)** |
|---|---|---|---|---|---|
| log Transcription rate, $\log(\alpha_m)$ | 2.764 | 2.784 | 2.782 | 0.054 | 0.052 |
| log Translation rate, $\log(\alpha_p)$ | 0.239 | 0.259 | 0.255 | 0.055 | 0.053 |
| Transcriptional delay, $\tau$ | 30.0 | 30.546 | 30.652 | 6.653 | 6.578 |

Table S3: The true values for the parameters which were used to generate the data in Figure 3A, alongside the means, $\boldsymbol{\mu}$, and standard deviations, $\boldsymbol{\sigma}$ of the corresponding one-dimensional posterior distributions, from both the MALA and MH algorithms (Figure S2).

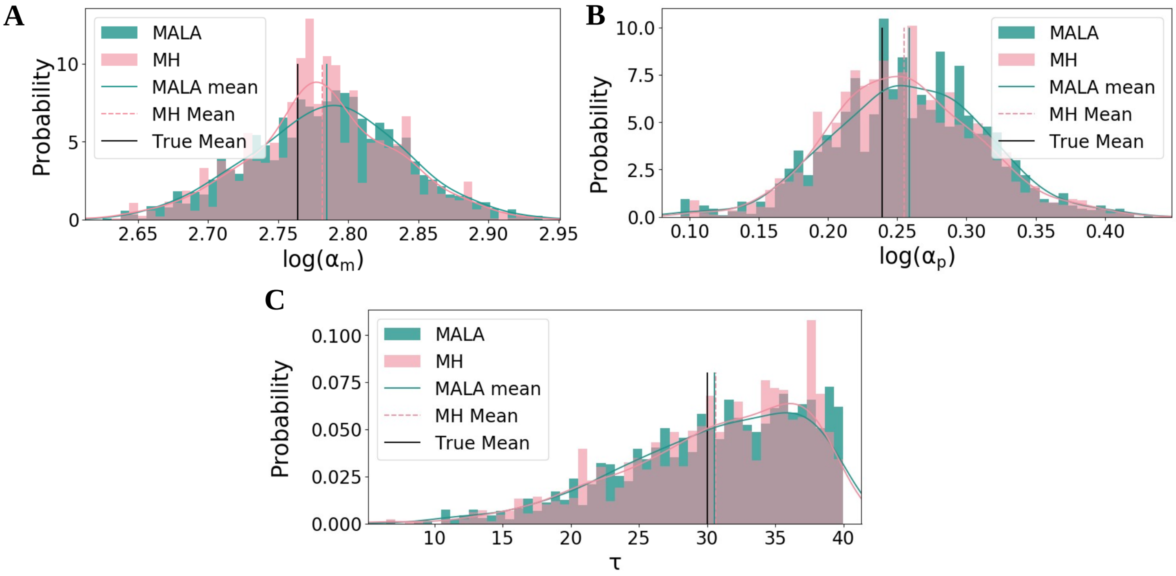


Figure S2: **Our algorithm accurately samples posterior distributions. A, B, C.** Histograms for both MALA and MH on the 1-dimensional problem for the transcription rate (A), translation rate (B) and transcriptional delay (C). Histograms and kernel density estimates are plotted, alongside the mean from each chain and the ground truth value. Other parameters are shown in Figure 2.

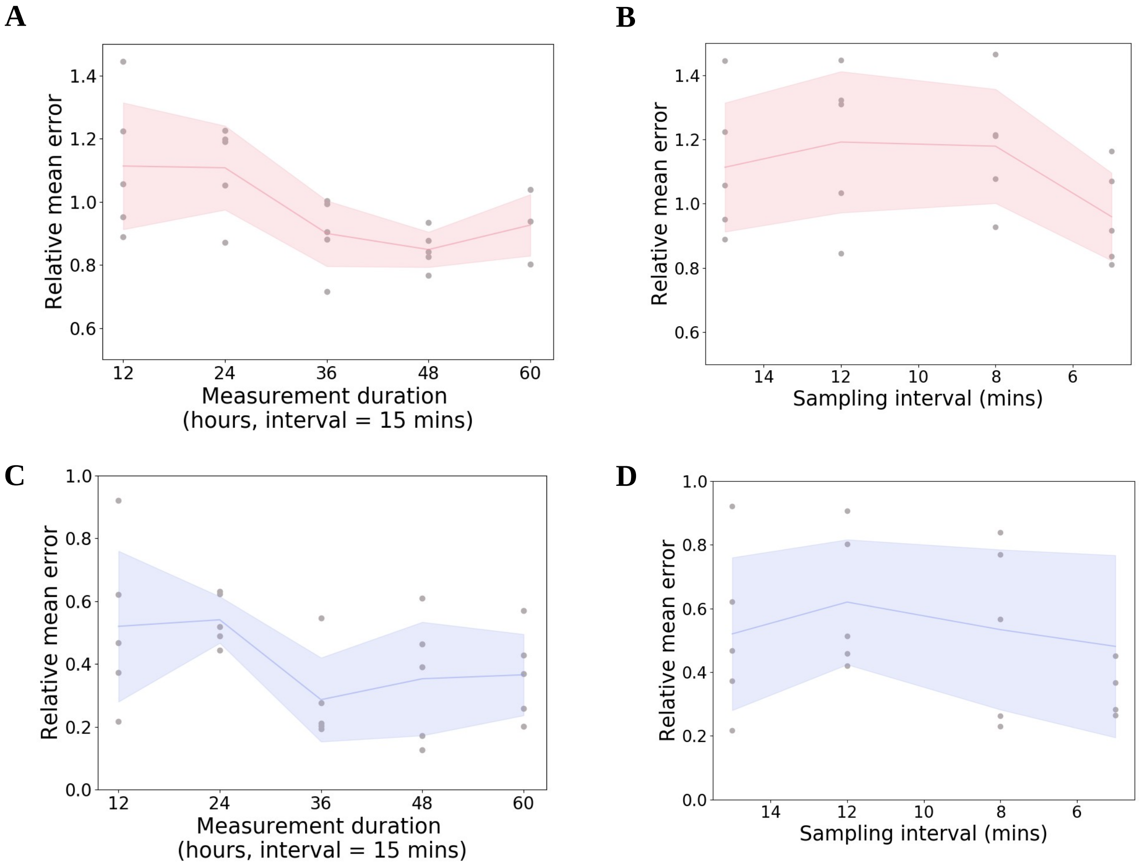


Figure S3: **Increasing the length of time course data improves inference more than increased sampling frequency.** **A.** $\mathrm{ME}_{\boldsymbol{\theta}}$ for low coherence data sets sampled with different lengths, from 12 hours to 60 hours. **B.** $\mathrm{ME}_{\boldsymbol{\theta}}$ for low coherence data sets sampled with different intervals. **C** Same as A with the high coherence data sets. **D.** Same as B with the high coherence data sets.

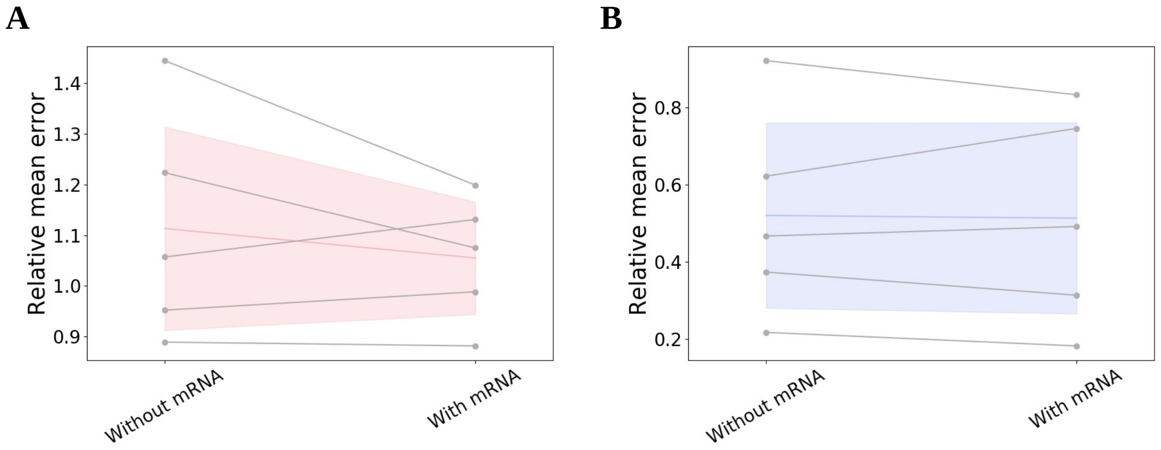


Figure S4: **Additional measurements of mRNA copy numbers do not improve overall inference accuracy. A.** Values of $\mathrm{ME}_{\boldsymbol{\theta}}$ (see equation (S36)) for low coherence data sets from Figure 5 with and without additional data on mRNA copy numbers (see supplementary Section S.9 for details). **B.** $\mathrm{ME}_{\boldsymbol{\theta}}$ for high coherence data sets from Figure 5 with and without additional data on mRNA copy numbers. The coloured lines and shaded areas represent the mean and standard deviation across observed values of $\mathrm{ME}_{\boldsymbol{\theta}}$, respectively.

**A**

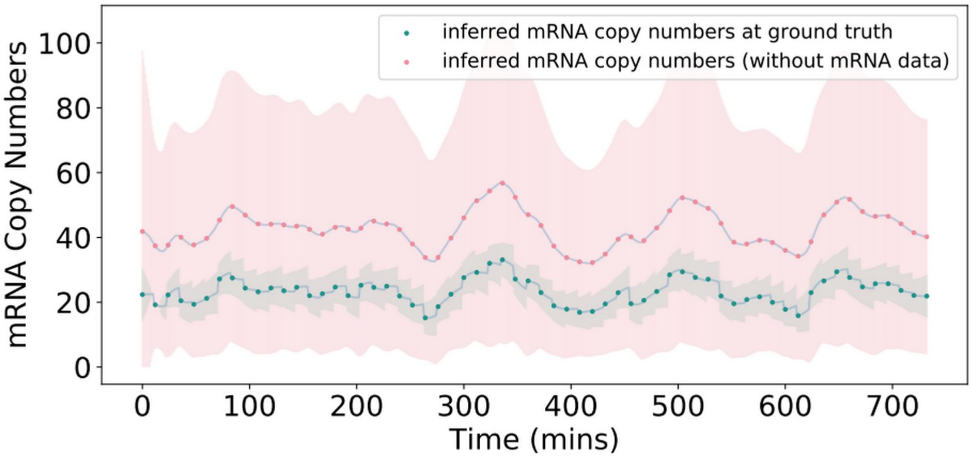
inferred mRNA copy numbers at ground truth
inferred mRNA copy numbers (without mRNA data)
mRNA Copy Numbers
0
20
40
60
80
100
0
100
200
300
400
500
600
700
Time (mins)

**B**

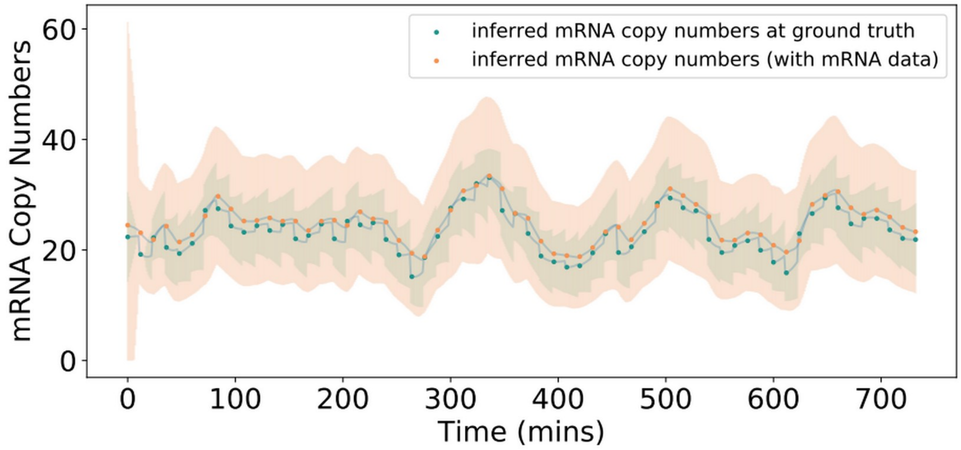
inferred mRNA copy numbers at ground truth
inferred mRNA copy numbers (with mRNA data)
mRNA Copy Numbers
0
20
40
60
0
100
200
300
400
500
600
700
Time (mins)

Figure S5: **Inclusion of data on mRNA copy numbers reduces uncertainty on estimated mRNA time series.** **A.** The distribution of posterior inferred mRNA copy numbers from the Kalman filter is shown (pink line: mean, pink shaded area: 95% confidence interval). The mean was calculated using mean inferred mRNA copy number across 1000 randomly chosen posterior samples at each time point, and the variance was calculated from the same samples using the law of total variance, thus accounting for uncertainty on the model parameters and for uncertainty on mRNA copy numbers at each posterior parameter combination. Additionally, we show the inferred mRNA copy numbers if only the ground truth parameter combination is considered in the Kalman filter (green line: inferred mean, green shaded area: 95% confidence interval). The posterior mean estimates are far from the ground truth levels, with the confidence interval covering values up to three times larger than the ground truth mean. Ground truth parameters are as in the high coherence parameter set of Table S2. **B.** If we include mRNA distribution information from an *in silico* smFISH experiment, the uncertainty on the inferred mRNA copy numbers is significantly reduced (orange dots: inferred mean, orange shading: 95% confidence interval), and also centered on the ground truth estimate (green dots and shading as in A).

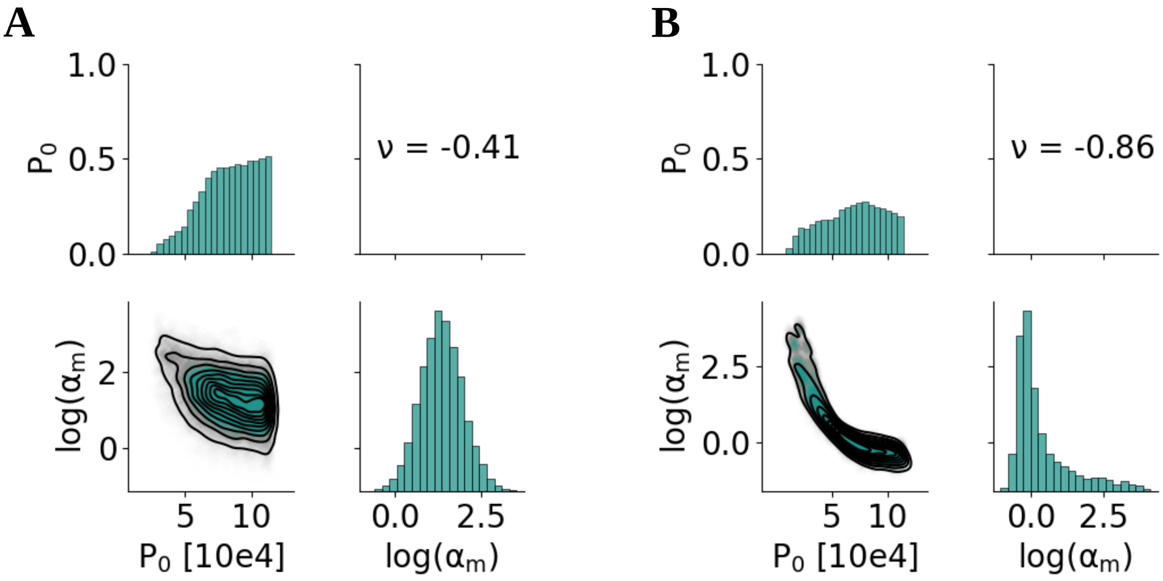


Figure S6: **Additional data on mRNA copy numbers constrains relationships between parameters.** **A.** Joint posterior distributions for the $P_0$ and $\log(\alpha_m)$ parameters, corresponding to the data set analysed in Figure 6A. The parameters are not highly correlated. **B.** Posterior distribution on the same data as in A and with mRNA distribution information included in the inference method. The relationship between the parameters is now more strongly defined. However, the support of the individual marginal posteriors remains large. This illustrates that the relative uncertainty $\mathrm{RU}_{\boldsymbol{\theta}}$ may not be reduced despite improvements in the estimation of $\alpha_T$ (Figure 6D).

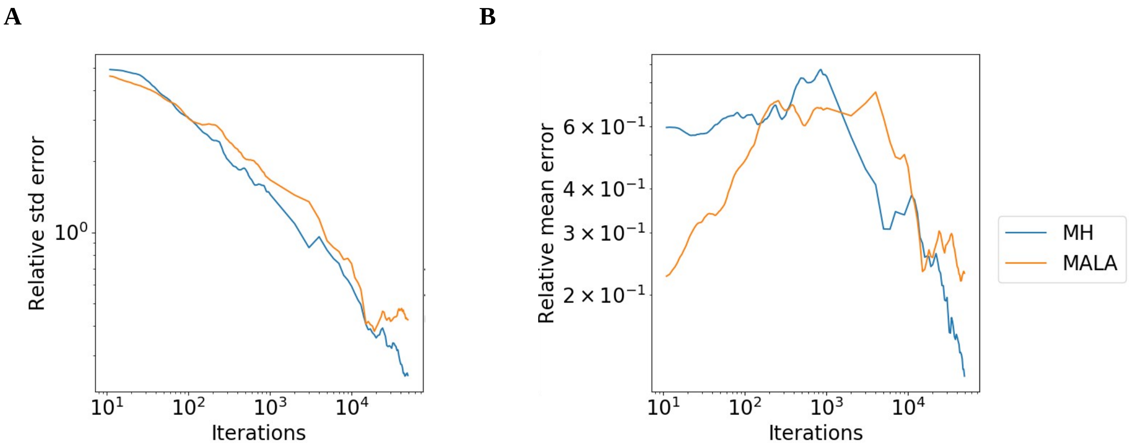


Figure S7: **MH and MALA converge to the true posterior distribution at a similar rate** Relative error in the **A** standard deviation and **B** mean of the posterior distribution as an increasing number of samples is drawn using MALA or MH, for an *in silico* data set of protein expression with a duration of 12 hours and with a sampling interval of 15 minutes, as well as assuming five unknown parameters. Note, that the data in Figures 3, 4 and 5 have the same structure. The *in silico* data set was generated using the parameter combination $P_0 = 47515, h = 4.77, \mu_m = \log(2)/30, \mu_p = \log(2)/90, \alpha_m = 2.65, \alpha_p = 17.61, \tau = 38.0, \Sigma_\epsilon = 10^6$. To calculate the error, a ground truth posterior distribution is generated by drawing 1,600,000 samples using MH. Errors are averaged over 8 chains for each sampler. For each chain, relative differences from the ground truth are calculated on each unknown parameter and then summed, to obtain an error estimate for the standard deviation and the mean. Before errors are calculated, chains are warmed up as described in supplementary Section S.3.
